# The dynamic role of HLA proteins on compositional alterations of T-cell repertoires in inflammatory bowel disease

**DOI:** 10.1101/2025.10.27.683825

**Authors:** Evgeniya Lokes, Gabriele Mayr, Malte Ziemann, Bernd Bokemeyer, Stefan Schreiber, Astrid Dempfle, Kazuyoshi Ishigaki, Andre Franke, Hesham ElAbd

## Abstract

**Background:** Inflammatory bowel disease (IBD), which is characterised by genetic predispositions and dysregulated immune responses, is rapidly emerging as a global health challenge. Genetic variations in the human leukocyte antigen (HLA) region are strongly associated with IBD; nonetheless, the functional consequences of this variation on the composition of T-cell receptors remain poorly understood.

**Methods:** We conducted comprehensive CDR3-QTL mapping using T-cell receptor beta (TRB) repertoires paired with HLA allotypes from 1,973 individuals, including 1,201 individuals with IBD and 772 healthy controls (HCs), to explore the role of the HLA allelic variants on TRB composition. Using network analyses, we defined key CDR3 motifs of public clones that were linked to risk HLA alleles for chronic inflammatory diseases.

**Results:** We identified novel sites within both HLA class I and class II proteins that were strongly linked to TRB amino acid composition – *cdr3*QTLs, in both HCs and individuals with IBD. Those sites in HLA-DRB1 and HLA-DQ had stronger effects on CDR3 composition than did the disease in the IBD cohort. In HCs, but not in UC or CD, the strongest HLA signals that affected expanded clones, overlapped with primary CD risk loci from GWAS, *e.g.*, DRB1 site 70 and DQA1 site 25. The strongest CD-specific effects on TRB composition were found in HLA-B, especially at sites that modulate viral responses (*e.g.*, HLA-B sites 9, 67). Finally, the main risk HLA alleles for chronic inflammatory diseases clustered together based on the physicochemical properties of residues mapped to *cdr3*QTLs, suggesting that risk alleles might exert similar effects on the TRB repertoire.

**Conclusion:** Structurally, the main *cdr3*QTLs in both HLA class I/II are located in peptide-binding sites or sites contacting TCRs, highlighting their direct and antigen-mediated influences on TRB repertoires. Our findings suggest that *cdr3*QTLs in HLA class I exert IBD-specific effects on the TRB composition, influencing the dysregulated T cell responses implicated in IBD pathogenesis, possibly on the earlier stages of T cell development. While HLA class II *cdr3*QTLs show universal effects and strong associations with T-cell receptors, irrespective of disease.

## Background

Inflammatory bowel disease (IBD) is undergoing a rapid transition from a rare condition to a global health challenge (1–3), with the prevalence in some regions now exceeding 1% (4,5). IBD is subdivided into Crohn’s disease (CD) and ulcerative colitis (UC), both characterised by chronic, relapsing inflammation of the gastrointestinal tract, in which T lymphocytes play a crucial role in driving dysregulated immune responses (6–12). Although environmental factors (13), such as the microbiome (14–17,13) and infections (18–24), contribute to IBD pathology, genetic susceptibility remains a key determinant. Among the identified loci, including *ATG16L* and *NOD2,* the human leukocyte antigen (*HLA*) locus stands out as one of the strongest and most consistent risk factors in multiple chronic inflammatory diseases, not only IBD (19,21). Classical HLA proteins are classified into class one (HLA-I: HLA-A, HLA-B, HLA-C) and class two (HLA-II: HLA-DR, HLA-DQ, HLA-DP); both classes are extremely polymorphic with thousands of known alleles, particularly for HLA-B and HLA-DRB1 (25). Recent advances have highlighted the role of HLA variants in modulating and selecting specific properties of T cell receptor (TCR) repertoires in healthy individuals (26–29), as well as during autoimmune conditions (30) and cancer (31).

Antigenic peptides are presented to T cells as extended helical fragments bound within a groove-shaped peptide-binding domain, which is made from two helices flanking an 8-stranded beta sheet. In HLA class I, the entire peptide binding domain is composed entirely from the variable alpha chain, while in HLA class II, it is assembled from two chains, *i.e.*, alpha and beta. Peptide binding specificity and affinity of the domain are determined by the physicochemical properties of the variable interaction pockets – peptide-binding sites (PBS), that accommodate five peptide core positions 1, 4, 6, 7 and 9, denoted A-F in HLA-I, P1-P9 in HLA-II. Other amino acids located on the surface of this domain are generally exposed for direct contact with the T cell receptor.

A diverse collection of TCRs is generated through V(D)J recombination, where variable (V), diversity (D), and joining (J) gene segments assemble to form the antigen-binding site. Within this structure, the <u>c</u>omplementarity-<u>d</u>etermining regions (CDRs) are arranged such that CDR1 and CDR2 predominantly contact the HLA molecule, whereas the highly diverse CDR3 loop primarily engages with peptides located in the HLA peptide-binding domain. The diverse CDR3s on the T-cell receptor β chain (TRB) enable T cells to recognise a wide range of peptide-HLA complexes.

In the current study, we aimed to identify HLA-specific effects that pregovern CDR3 repertoires involved in IBD pathogenesis. For that, we conducted comprehensive CDR3-QTL mapping (27), employing high-throughput bulk TCR sequencing matched with HLA genotyping in a large cohort of 1,973 individuals with and without IBD. We hypothesised that HLA class I and class II variants play a dynamic role in the immune repertoire, given that multiple layers of HLA-TCR interactions occur during different stages of T cell development. We also aimed to elucidate the functional consequences of the main HLA risk alleles for chronic inflammatory diseases by identifying the CDR3 signatures of public clones shaped by these alleles.

## Methods

### Cohorts and sample collection

For the analysis, we used previously published datasets of TCR-beta chain (TRB) immune sequencing data from 772 healthy blood donors (32) and 1,201 IBD patients (12): 896 individuals with CD and 305 individuals with UC. The TRB datasets were deeply sequenced, with mean repertoire sizes X̄_CD_ = 208,066 unique CDR3β sequences, X̄_UC_ = 239,989 unique CDR3βs, and X̄_HC_ = 239,146 unique CDR3β clonotypes. Our analysis focused on productive TRB sequences, which composed ∼83% of the overall repertoire of healthy individuals and those with IBD. We excluded 52 individuals with IBD from the subsequent analyses, as their repertoires consisted of fewer than 30,000 clonotypes.

### TCR repertoire profiling

T-cell receptor (TCR) beta chain repertoires were profiled via the Adaptive Biotechnologies immunoSEQ® Assay (Adaptive Biotechnologies, Seattle, WA, USA). Genomic DNA was extracted from peripheral blood mononuclear cells (PBMCs) via a DNeasy Blood Extraction Kit (Qiagen). Up to 18ug of DNA was used to profile the repertoire via the immuneSEQ assay using a biased controlled multiplex PCR reaction (33). After amplification, samples were pooled together and sequenced using high throughput next generation sequencing (34). After sequencing and demultiplexing, clonotypes were identified and their expansion was quantified from the generated sequencing reads. Subsequent bioinformatics analysis was performed with custom scripts to assess the clonality, diversity, CDR3 amino acid composition, shared CDR3 motifs of public clones, as were CDR3-QTL analyses.

### HLA imputation from genotyping chips

Human leukocyte antigen (HLA) alleles were imputed from genome-wide genotyping data. Genotyping was performed via the Illumina Global Screening Array. Prior to imputation, standard quality control (QC) procedures were applied to the genotype data, including the removal of samples with high missing call rates >5%, low genotyping quality D-scores < 0.9, and low call rates <95%. HLA allele imputation was performed using our inhouse imputation pipeline (https://github.com/ikmb/hla), which rely on a comprehensive, high-resolution multiethnic reference panel (35). Following imputation, only HLA alleles with an imputation quality exceeding a predefined threshold > 0.9 were retained for downstream analyses. The imputed HLA alleles were then represented at 4-digit resolution for the classical HLA-A, HLA-B, HLA-C, HLA-DRB1, HLA-DQA1, and HLA-DQB1 loci.

### HLA preprocessing

For each HLA allele, we acquired HLA protein sequences from IMGT (https://www.ebi.ac.uk/ipd/imgt/hla/ dated 03.05.2024). By comparing amino acid sequences of HLA molecules at the mature protein level, we identified variable sites with at least two amino acid differences across alleles present in both datasets; for example, HLA-DQA1 at site 130 has either a polar hydrophilic serine (S) or a nonpolar hydrophobic alanine (A). For each variable HLA site, we built a matrix of site-variant carriages with individuals as rows and observed amino acids as columns, where each individual had either 0, 1, or 2 occurrence values for each amino acid variant (**Figure 1A**). We considered HLA sites to be monomorphic if 95% of site variants gave the same amino acid at this site, those sites were removed from the analysis (meaning the site is not variable when 95% of alleles had the same amino acid at this site), and from analyses, we removed all site variants that were present in less than 5% of individuals.

### CDR3 normalisation

We included only productive CDR3 sequences with lengths ranging from 12 to 18 amino acids and analysed their amino acid composition at each position, according to the IMGT nomenclature (36), to account for the site-specific effect of HLA on CDR3β. We approximated *“naïve”* repertoires computationally by collapsing existing clonal expansions into singletons, which allowed us to specifically investigate processes during thymic selection rather than those in the periphery. CDR3β sequences were mapped to the IMGT positions, with a focus on the middle CDR3β positions 106-118, which directly interact with the peptide-HLA (pHLA) complex. In all analyses, if not explicitly stated, we analysed the CDR3β composition at the level of amino acid frequency at each CDR3β position, accounting for the CDR3β length. CDR3β amino acid frequencies at each IMGT position were calculated after excluding germline-encoded regions derived from the V and J segments. When V or J segments were not resolved to the gene level for the CDR3β sequence, we removed all amino acids common to the V or J family from the CDR3β sequence.

We used inverse rank transformation (IVT) for the CDR3β amino acid frequencies across CDR3β positions at each CDR3β length to normalise the distribution of CDR3 amino acid frequencies, as the normal distribution of frequencies is a prerequisite for multivariate multiple linear regression (MMLR), described in the following section. IVT was performed by ranking the CDR3 amino acid frequencies between patients and scaling the ranks back to the original scale. Rare CDR3 amino acids, which were observed at each CDR3β position in less than 50% of individuals, were excluded from the CDR3-QTL analysis because of the violation of model assumptions.

### Public CDR3 clones linked to HLA alleles

To identify potential associations between specific HLA alleles and the presence of public TRB clonotypes, a series of two-tailed Fisher’s exact tests were performed. Public TRB clonotypes were defined as those identified in at least 5% of unrelated individuals within the combined healthy and IBD cohort. The choice of 5% threshold for a public clone was chosen to match the minor allele frequency (MAF) for HLA alleles. After reducing the threshold for a public clone to be present in at least two repertoires, individuals shared more than 14,2% productive TRB sequences.

For each unique combination of an HLA allele (at 4-digit resolution) and a public TRB clonotype, a 2ξ2 contingency table was constructed. This table categorises individuals on the basis of homozygosity, heterozygosity and absence of the specific HLA allele and presence or absence of the specific public TRB clonotype. Given the substantial number of tests conducted (approximately 1.3ξ10^9^ tests derived from 40 common HLA alleles ξ 32 ξ 10^6^ public TRB clonotypes), a stringent multiple testing correction was applied. An adjusted P value (Bonferroni-corrected p value) threshold of p < 0.05 was considered statistically significant. All the statistical analyses were performed via custom scripts using R statistical software (version 4.3.2).

### CDR3-QTLs with multivariate multiple linear regression

We tested associations between normalised CDR3 amino acid frequencies for each CDR3β length‒position combination and variable HLA sites using the CDR3-QTL approach (27), which uses multivariate multiple linear regression (MMLR). In this work, we refer to HLA amino acids as “sites” and CDR3β amino acids as “positions” to avoid confusion. We used MMLR to model the relationships between the amino acid variants at each variable HLA site as independent variables and the CDR3 amino acid frequency at each CDR3β position for each CDR3β length as a compositional matrix of dependent variables. We used the following full model:

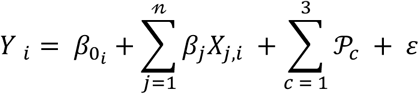

where 𝑌*_i_* is a matrix of CDR3 amino acid frequencies at CDR3β length-position *i,* 𝑋*_j,i_* is a matrix of amino acid occurrence at a variable HLA site *j* with *n* amino acid alleles, 𝒫*_c_* is a matrix of principal components (PCs) *c* of all patients’ HLA alleles, 𝛽_0_ represents the intercept and 𝜀 represents the error.

For the null model, we used the principal components of all HLA alleles as a set of independent variables:

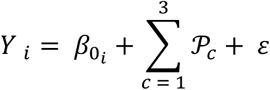

We assessed the significance between the variance of the full and null models via MANOVA with Pillai’s trace. We used the first nine PCs as covariates in the full and null models and removed correlated HLA alleles (r^2^>0.5) before calculating the PCs. We defined the optimal number of PCs on the basis of the permutation test, where we compared the cumulative variance explained by the PCs for the shuffled HLA alleles with corresponding patients’ IDs (n _permutations_ = 1000), which allowed us to identify that nine PCs capture a significant genotype structure, as the p value for the additional tenth PC was above the significance threshold.

### Variance partitioning framework

To disentangle the contributions of HLA variation and disease group status to TCR repertoire composition, we implemented a variance partitioning framework on the basis of nested multivariate multiple linear models. To test whether the impact of HLA on the CDR3β composition is stronger than that of disease effects, we trained four different CDR3-QTL models to explain the CDR3β composition by providing HLA genotypes together with corresponding sample phenotypes: HC, CD or UC. We compared the effects of HLAs, which are conditioned on phenotypes with the effects of phenotypes, with those of HLAs. Let *CDR3* denote the response matrix of CDR3 amino acid frequencies at each tested nontemplate position (according to IMGT), *X_HLA_* denote the matrix of HLA predictors, *groups* the categorical disease status, and PCs denote the principal components used as covariates. We fitted the following four models:

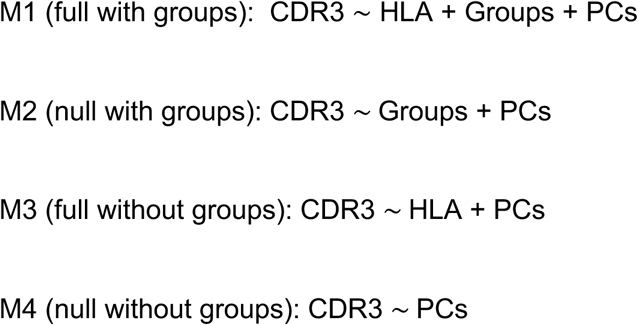

The significance of variance explained by HLA variation and disease groups was assessed via omnibus tests implemented in the mmvm package (R). For each predictor set, we fitted multivariate models of CDR3 composition and obtained p values from the omnibus test, which evaluates the null hypothesis that the corresponding predictor set jointly explains no additional variance in the multivariate outcome. The reported p values, therefore, reflect the overall multivariate contribution of HA site variants or groups, conditional on the covariates included in each model. We applied the following conditioning strategies:

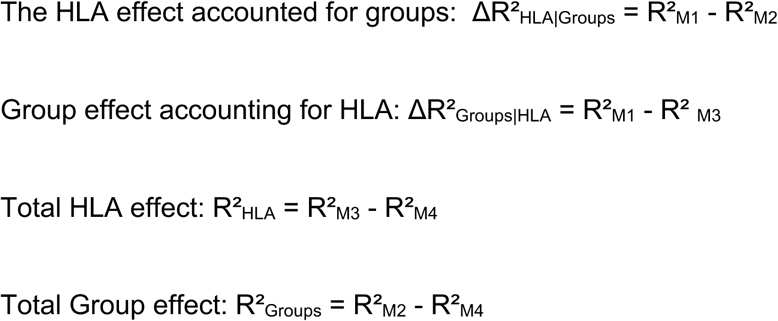

Partial R^2^ values were additionally computed to express the effect sizes of the HLA and groups conditional on each other. The significance of the incremental variance explained was assessed via MANOVA Pillai’s trace between nested models.

### Expansion-weighted downsampling of TRB repertoires

To harmonise sequencing depth while preserving relative clonal expansions, we downsampled each repertoire to a common target depth 𝑆 using probability-proportional-to-expansion sampling. This kept highly expanded clones more likely to be retained than rare ones, opposite to the random subsampling of unique CDR3s (**Figure 3A**). We employed a multifaceted approach to investigate HLA associations with TRB amino acid composition across different T cell activation states. First, we assessed the effects of HLA on the composition of thymically selected clones by collapsing all clonally expanded sequences into singletons and treating all the clones as unique sequences. Second, to understand how HLA influences the result of peripheral activation, we examined its association with the composition of primed and expanded clones, calculating CDR3 amino acid frequencies accounting for the expansion. Finally, to bridge these insights, we also investigated HLA associations with the composition of clones that were highly expanded but were analysed in their preexpansion (*expanded singletons*) state, providing insight into early determinants of an antigen-driven response. We used two different approaches for the CDR3 downsampling of expanded clones: first, when TCR clone expansion increased the chance for the TCR clone to be included in the down ampled TCR repertoire; second, we used a random selection of the desired repertoire size.

### Identifying unique sites via step-forward conditional *cis-*haplotype analyses

Conditional *cis-*haplotype analysis with a step-forward approach was used to disentangle relationships between statistically significant but genetically linked HLA sites (27,37) by excluding the influence of genetic linkage and the high correlation between significant sites within each HLA protein. We followed the procedure described in previous studies (27,37). Briefly, after the initial CDR3-QTL analyses, each HLA gene was analysed as follows: the most statistically significant *cdr3*QTL signal was serially added as a covariate to the step-forward regression testing the remaining HLA sites within the region until the improved model achieved saturation, and no statistically significant signals were further observed. Each round of conditional *cis-*haplotype analysis transformed the initial HLA occurrence matrix of independent variables into a new matrix of independent variables with the HLA haplotype occurrence matrix, containing the tested HLA site and the strongest *cdr3*QTL signal from the previous round. We performed conditional haplotype analyses on combined healthy and IBD datasets, as well as on stratified into HC, CD and UC to see whether the disease confounds the set of independent *cdr3*QTLs.

### PERMANOVA and permutation analysis

Permutational multivariate analysis of variance (PERMANOVA) was performed to test *cdr3*QTL signals identified by MANOVA. Robust Aitchison transformation for CDR3 frequency was used, and each matrix of amino acid sequences for each IMGT position and CDR3 length was regressed as a function of HLA genotype. We used PERMANOVA as a nonparametric technique, which does not assume normality or homogeneity of variances but accounts for the compositional nature of the TCR-seq data.

### Power analysis

The power analysis was performed by serially downsampling the sample size for cohorts, starting with 400 samples, gradually increasing the sample size and performing the CDR3-QTL independently with each sample size. A larger sample size increases the statistical power to discover novel HLA sites outside the HLA-DRB1 region that act as *cdr3*QTLs.

### Code availability

The scripts supporting the analyses used in this study are available at https://github.com/ikmb/CDR3_QTL.

## Results

### Study design and overview of TRB and HLA genotyping datasets

We utilised TCRβ (TRB) sequencing data of rearranged CDR3 regions from two distinct cohorts (**Figure 1A**): healthy controls (HC, n = 772) (32) and individuals with IBD (IBD, n = 1,201: 896 individuals with Crohn’s disease (CD) and 305 individuals with ulcerative colitis (UC)) (12). TRB repertoires were paired with HLA class I and II allotypes, imputed from high-density genotyping arrays (**Figure 1A, S1, S16**). We were interested in pinpointing disease-specific influences of HLA proteins on the TRB repertoires in the context of IBD. Unless otherwise specified, we focused on productive TRB sequences, which composed ∼83% of TRB clonotypes regardless of phenotype (**Figure S2B**).

We asked two main questions: (Q1) which TRB repertoire conditions and underlying mechanisms best reflect IBD-specific HLA effects, and (Q2) whether IBD HLA risk loci are mechanistically involved in shaping immune repertoires at the level of amino acid composition. In our primary discovery analyses, we addressed three conditions to answer Q1 (**Figure 1A**) regarding the influence of HLA on TRB repertoires: (i) HLA effects on TRB singletons, mimicking *naïve* repertoires; (ii) HLA effects on TRB repertoire, while accounting for the clonal expansion; and (iii) HLA associations with the composition of highly expanded clones without considering their expansion, referred to as *expanded singletons*. We hypothesised that HLA effects on *expanded singletons* will be limited to HLA class I, given a massive expansion of antigen-experienced CD8^+^ relative to CD4^+^ T cells (38), while HLA effects on the singletons and TRB clones considering their expansion would be more prominent in HLA class II, given the overall abundance of CD4^+^ T cells in blood samples. To answer these questions, we employed distinct computational strategies, including TCR expansion-weighted selection (**Methods**).

**Figure 1.**
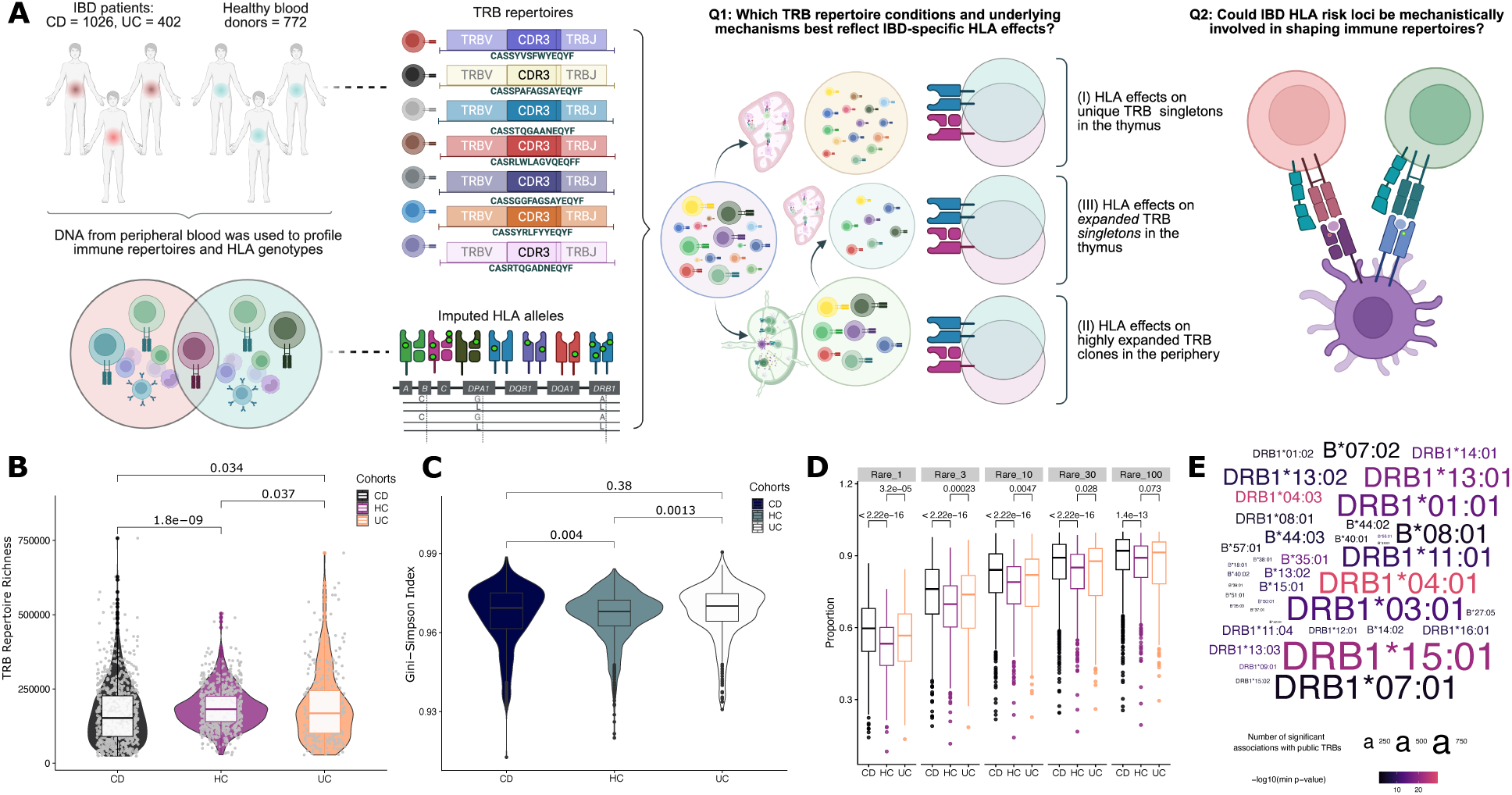
Study design and overview of TRB repertoires in HC and IBD cohorts. **(A)** DNA from the peripheral blood of the IBD and HC cohorts was used to profile TRB CDR3 repertoires and impute HLA alleles; HLA amino acid sequences were profiled from imputed HLA alleles and transformed into variable sites to focus on site-specific effects (marked as green dots). We used a multifaceted approach to investigate HLA effects on TRB amino acid composition across different T-cell developmental stages to answer Q1. In a yellow circle, HLA affects the composition of CDR3 singletons. In a green circle, HLA influences on expanded clones. In a pale blue circle, HLA associations with the composition of highly expanded clones in their pre-expansion state (*expanded singletons*), providing insight into early determinants of an antigen-driven response. Given known IBD risk loci from GWAS studies (*20,21,39*), we were interested whether they play a role in shaping immune repertoires. **(B-D)** Overview of the TRB repertoires of the cohorts. **(B)** TRB repertoire richness, pairwise group pairwise comparison with ANOVA P = 3.93e-07, Tukey multiple comparison tests: PHC-CD = 1.8e-09, PUC-CD = 3.4e-02, PUC-HC = 3.7e-02. **(C)** Gini‒Simpson diversity index, a comparison between groups. Compared with HCs, patients with CD and UC presented greater median diversity (PHC-CD = 4e-03, PUC-HC = 1.3e-03, PUC-CD = 0.38), even though the lower tails tended to be longer, indicating increased heterogeneity in diversity and clonality among individuals with IBD. **(D)** Group pairwise comparison of the proportion of rare clones within immune repertoires, starting with unique clones (each dot represents the proportion of rare clones within each sample), until the expansion of rare clones reached 100. We observed a significantly increased proportion of rare clones in the IBD cohort across all fractions of rare clones, except for the last comparison of 100 rare clones between HCs and UCs. **(E)** HLA-B and HLA-DRB1 alleles, associated with TRB clones, the font size stays for the number of associated clones, and colour shows the p value of the strongest association with certain HLA allele.

### High heterogeneity without global clonal expansion of TRB repertoires in IBD blood samples

While the TRB repertoire of HCs presented the highest median repertoire richness, significantly surpassing that of both CD patients (p=1.8 ×10^-9^) and UC patients (p=0.034) (**Figure 1B, S2A**), some individuals within disease cohorts presented exceptionally high richness, potentially representing unique immune states. Compared with that of healthy controls, the Gini–Simpson index, reflecting repertoire diversity and evenness (with higher values indicating a greater probability that two randomly selected sequences belong to different clones), was elevated in both the CD and UC repertoires (**Figure 1C**), in contrast to a previous report (40). We observed reduced clonality within the top 10 to 10^4^ clones between CD patients and HCs (**Figure S2D**), together with an increased proportion of rare clones in IBD patients (**Figure 1D, S2F**), indicating more even and heterogeneous TCR repertoires in peripheral blood, with fewer dominant clonal expansions than in healthy controls. The contribution of the top 100,000 expanded clones was greater, reflecting a repertoire structure with fewer dominant clones and an expanded tail of low-frequency clones, which is consistent with increased peripheral T-cell diversity, as observed in a recent study on IBD-discordant monozygotic twins (9). Thus, broadly heterogeneous repertoires provide an ideal context to detect HLA effects on TRB sequences, as subtle HLA site-specific effects (**Figure S7**) are less likely to be masked by large clonal expansions.

### HLA alleles leave detectable signatures on public TRB clones

Previous studies have demonstrated strong relationships between certain HLA alleles and public TRB clones (41,42), including biased V and J usage (26,43). We proposed that HLA molecules would imprint detectable sequence signatures on TCR repertoires. As a critical proof-of-concept, we screened the most variable, nonetheless common HLA alleles in HLA-DRB1 and HLA-B. Among 90 alleles found in both cohorts, 39 HLA alleles we found to be strongly linked to 7,262 public TRB clone (Fisher’s exact test P_FDR_<0.05 after false discovery rate (FDR) correction, or 277 clones with p_adj_<0.05 after Bonferroni correction) (**Figures 1E, S4**) (**Methods**). The strongest association was observed between HLA-DRB1*04:01 and the TRB clone CASSEASGGADTQYF (TRBV6-1, TRBJ2-3) (p_FDR_<1.15×10^-29^), with currently unknown antigen specificity. A public TRB clone was defined as being present in at least 5% of individuals to match minor allele frequency of common HLA alleles **(Methods)**. The total number of 10,876 public clones were identified, representing no more than 0,013% of all productive TRB sequences (**Figure S3)**.

We found that public TRB clones in both the IBD and HC cohorts that were strongly linked to HLA alleles, shared common CDR3 motifs (**Figure S4**). We compared Aitchison distances between public TRB clones that were strongly associated with HLA alleles, and those displaying no association with HLA. Most clonotypes were restricted to HLA-II alleles, corroborating previous observations by us (41) and others (42,44,45). We found that the clones linked to HLA alleles exhibited distinct compositional signatures at the highly variable middle positions of the CDR3, unlike public clones without HLA association, on the example of HLA-DRB1*15:01 as it had the largest number of linked to it public clones and is a risk allele to IBD in European population (21) **(Figure S5)**. Nonetheless, the overall influence of HLA was more subtle than reshaping entire repertoires, acting instead in a position-specific manner on CDR3 (**Figures S7, S8)**. Given that HLA alleles vary in both the extent of their influence and in the presence of different residues at HLA protein sites, these strong HLA–TRB links provided the basis for extending our analysis to site-specific CDR3-QTL mapping (27) across entire repertoires (**Figure 2A**).

### Specific HLA class II sites have a stronger impact on the immune repertoire when accounting for disease

Previously, Ishigaki *et al.* revealed the role of HLA-DRB1 site 13 as a locus strongly affecting the composition of the CDR3 middle position in healthy individuals, namely, CDR3 quantitative trait loci, or *cdr3*QTLs (27). Inspired by this finding, we further investigated site-specific HLA effects on TRB repertoires at the level of CDR3 amino acid composition using a larger cohort of both healthy individuals and those affected by IBD. In essence, our primary CDR3-QTL discovery analysis systemically linked HLA site-specific allelic variants with CDR3β amino acid composition (**Figure 2A**) while accounting for other confounders, such as disease itself, by conditioning on IBD subtypes (**Figures 2C-D**). In the current study we use terms CDR3 and CDR3β interchangeably, since we had profiled TRB repertoires.

As a proof of concept, we validated previously reported *cdr3*QTL associations within HLA class I and II region in an independent population of healthy individuals (27) (**Figure S10**). First, we focused on Q1 TRB condition (i) (**Figure 1A**) by collapsing the entire repertoire to a set of CDR3 singletons, computationally mimicking *naïve* repertoires (**Methods**). We trained the model to explain the amino acid composition of highly variable middle positions of CDR3β as a function of HLA allotypes, combining both datasets (**Figure 2A**). Through serial downsampling, we confirmed that the available cohort sizes provided sufficient power to robustly detect site-specific effects of HLA allelic variants on CDR3 (**Figure 2D, S9**). A total of 9,533 statistically significant associations, namely, *cdr3*QTLs, were identified (MANOVA, Pillai’s trace P value < 4.6×10^⁻7^ = 0.01/26,880 total tests) (**Figure 2D**). Among these, 70.8% of *cdr3*QTLs included HLA class II protein sites, whereas 15.9% of *cdr3*QTLs linked HLA sites to one highly diverse middle CDR3β amino acid position 109, followed by positions 108, 110, and 113, accounting for more than 50% of all significant associations (**Figure S11A**).

**Figure 2.**
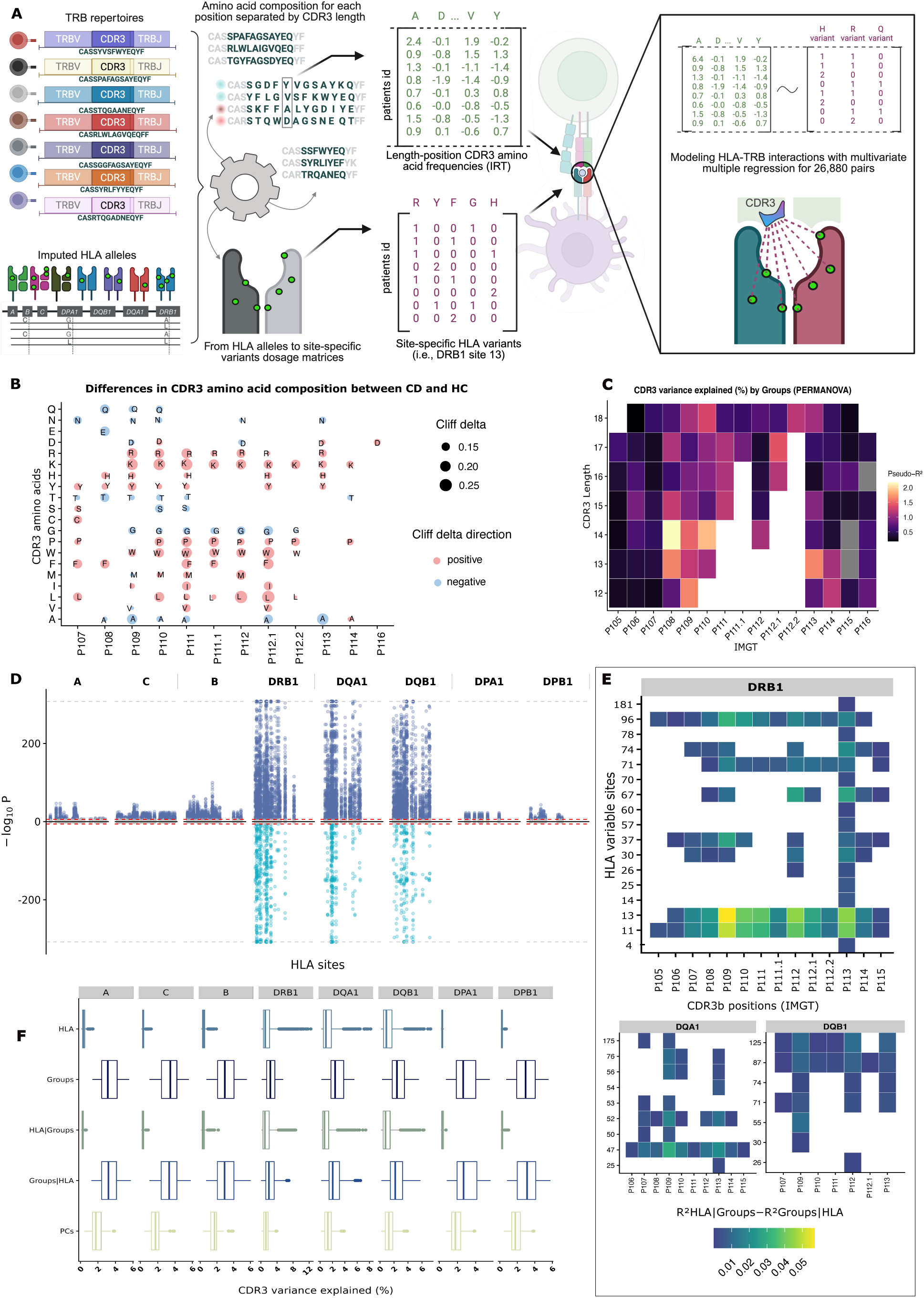
CDR3-QTL discovery analyses of combined IBD and HC datasets. **(A)** CDR3-QTL computational workflow. The CDR3 sequences, treated as unique sequences regardless their expansion, were sorted by their length; the amino acid composition for each CDR3 position was analysed separately. CDR3 amino acid frequencies were built from CDR3 sequences and inverted rank normalised. A total of 26,880 pairs of multidimensional matrices with CDR3 compositions (per CDR3 position and length) and HLA site variants were modelled with MMR (**Methods**), linking HLA variants with CDR3 composition. **(B)** The CDR3 amino acid composition differed between cohorts. CDR3 composition distributions were the most significantly different between CD patients and healthy controls (HCs), as tested at the position level (according to IMGT nomenclature) with PERMANOVA, and at the level of specific amino acids with the Mann‒Whitney U test: the Cliff delta test was used as a nonparametric estimation of correlations between amino acid frequencies and groups. **(C)** Preudo-R^2^ (PERMANOVA) of group status - phenotypes, on average, explained no more than 2.2% of the variation in the CDR3 composition. **(D)** Our results provide strong evidence that multiple HLA sites, across both Class I and Class II, significantly influence the CDR3β composition, with HLA sites on the X-axis and the significance of HLA‒ CDR3 associations (Pillai’s trace MANOVA P values) on the Y-axis; Miami plot illustrating the location of HLA sites (X axis) and the strength of associations between HLA and CDR3 composition (Pillai’s trace MANOVA P values) comparing all variable sites (upper part of the plot) with those showing the strongest effects of CDR3 regardless of disease (lower part of the plot). **(E)** While groups generally accounted for more variance via MANOVA, several HLA sites (Y axis) conditioned on groups showed markedly stronger effects than groups conditioned on HLA for explaining CDR3 composition in position-wise manner (X axis: P105-P115, IMGT nomenclature). **(F)** Differences in multivariate (R^2^) variance explained between HLA sites accounted for group effects and groups, accounting for HLA in the CDR3 composition. On average, Groups and Groups, accounted for HLA (Groups|HLA), explained a greater proportion of CDR3 variance than majority of HLA sites, with some drastic exceptions for specific HLA-DRB1 and HLA-DQ sites that explained CDR3 composition of combined cohorts better that groups.

We hypothesised that both HLA allotypes and disease affect the CDR3 composition (**Figures 2B-C**). To test this hypothesis, we considered four different models on combined TRB dataset to infer the mechanisms by which HLA sites, independent of disease, influence the CDR3 composition (**Methods**). We found that, on average, phenotype/groups (CD, CD, HC), accounted for HLA effects (Groups|HLA), had greater explanatory power for explaining the CDR3 composition than did majority of HLA class I and II sites (**Figures 2F, S11B**): 1.6% compared with 0.2%, respectively. However, several specific HLA-DRB1 and HLA-DQ sites, before and after accounting for group effects showed markedly stronger explanatory power to explained CDR3 composition for both cohorts, with, on average, ∼3.7% greater variance explained than did groups, accounted for HLA effects (**Figures 2E-F, S11A**). These site-specific effects highlight that global analysis may mask important HLA-driven signals, underscoring the need to stratify datasets to uncover disease-specific HLA influences on the TCR repertoire, even at the cost of reduced statistical power (**Figure S9**).

### *cdr3*QTLs in HLA-B shape both *naïve* and clonally expanded TCRs

To address Q1 conditions (ii) and (iii) (**Figure 1A**), we modelled diseases separately and analysed *cdr3*QTLs in segregated cohorts (CD, UC, and HC), employing computational strategies to model HLA-TCR interactions during different stages of T-cell development (**Figures 1A, 3A**). Importantly, we found that TRB sequencing depth influences the captured HLA effects on TRB repertoires, owing to the potential overrepresentation of highly expanded CDR3s at shallower depths (**Figures S12, S13**). Therefore, to ensure the robustness of our findings regarding the effects of HLA class I and II on the CDR3 composition, and to precisely distinguish HLA influences across *naïve* (singletons) and antigen-experienced (expanded) clones, and those at the stage before expansion (*expanded singletons*), in the following analyses we enriched immune repertoires with expanded clones using expansion-weighted downsampling (**Figure 3A, Methods**). Briefly, expansion-weighted selection of TRB clones was based on a clone’s proportional probability of inclusion to its original expansion, effectively enriching with highly expanded clones. For *expanded singletons* in the Q1 condition (iii) (**Figure 1A**), CDR3 amino acid composition was estimated without considering their expansion. Fixing TRB repertoire depth to a fixed numbers of 30,000 clones allowed us to assess the influence of HLA allelic variants on the composition of highly expanded TRB clones and *expanded singletons* consistently, independent of sequencing depth artifacts.

**Figure 3.**
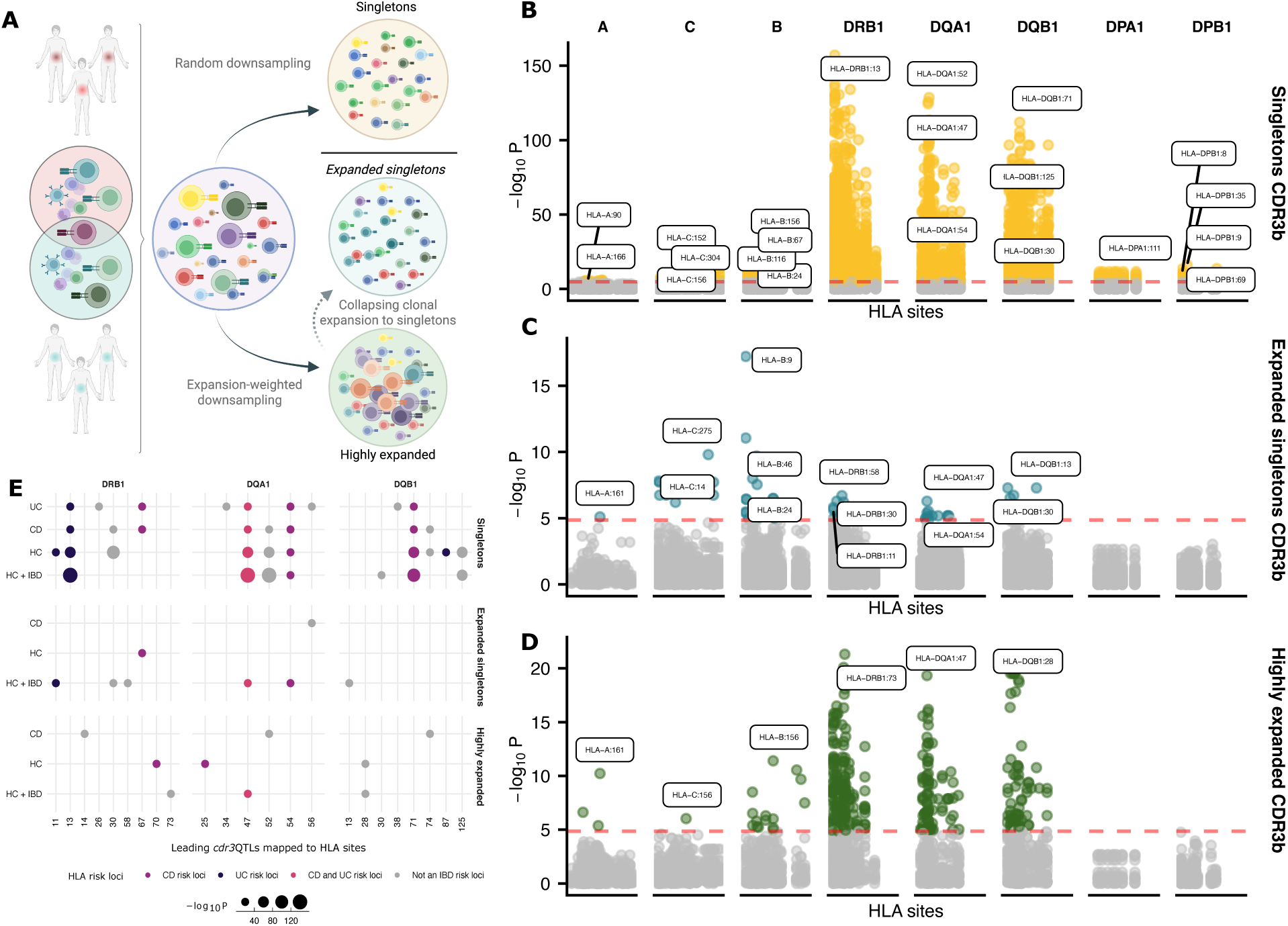
Expansion-weighted downsampling revealed differential effects of HLA class I from class II on T-cell singletons and expanded clones. **(A)** TRB repertoire downsampling strategies represent conditions stated in Q1: random downsampling for unique singletons in yellow, and expansion-weighted downsampling for *expanded singletons* in pale blue, and highly expanded in green: the clonal expansion increased the chance for the clone to be selected in the downsampled TRB repertoire, and the CDR3 amino acid frequencies were recalculated, based on the condition (**Methods**). **(B-D)** Manhattan plots with *cdr3*QTLs on the downsampled repertoire (size of 30,000) of (**B**) CDR3 singletons (yellow), (**C**) *expanded singletons* (pale blue), where leading *cdr3*QTLs in expanded singletons are HLA-B sites 9 and 46; and (**D**) highly expanded (green). The role of HLA Class I at expanded singletons, is the strongest. The MANOVA omnibus test was applied to the CDR3-QTL models to estimate the significance of associations between HLA variants at each variable site and the CDR3 amino acid composition. The red dotted lines in all the plots represent the stringent threshold for statistical significance according to the Bonferroni correction. **(E)** Overlap between *cdr3*QTLs on singletons, expanded singletons and highly expanded clones with HLA risk loci associated with CD and UC from Goyette et al.(39)

We hypothesised, that Q1 condition (ii) with expanded clones would reflect HLA effects on CD4^+^ T cell biology, given their relative abundance in the peripheral blood, but in the case of condition (iii) with *expanded singletons,* we expected it to reflect HLA effects on, predominantly, CD8^+^ T cells, given their relative expansion compared to CD4^+^ T cells. Our initial observations revealed striking differences in how and where HLA sites influence CDR3 compositions, also capturing different *cdr3*QTL patterns across disease groups, supporting our hypotheses (**Figure 3B**).

We saw earlier that HLA class II site variants, predominantly HLA-DRB1 and HLA-DQ, have an effect on *naïve* repertoires (**Figures 2D, 3B**). In contrast, when amino acid composition of antigen-experienced, hence expanded clones were calculated in the way to capture CDR3 characteristics before their expansion (i.e., *expanded singleton* repertoires) (**Methods**), HLA influences were observed primarily in HLA-B, with leading signals at sites 9, 46 and 67 (**Figures 3B, D**). Among 280 variable HLA sites in both class I and class II, except three sites in DRB1 and five in HLA-DQ, we observed a total loss of *cdr3*QTL signals within HLA class II on expanded CDR3 repertoires at the stage before expansion, supporting our hypothesis regarding capturing HLA effects on CDR3 composition of CD8^+^ T cells, which form complexes with HLA class I (**Figures 3B, D**). The more we enrich for highly expanded clones, the more pronounced was the HLA-B effect (**Figures 3D, S12B, S13**). Notably, HLA-B shaped CDR3 composition of expanded clones through sites that overlapped with sites that are directly involved in modulating responses to viral infections (*e.g.,* sites 9, 67, and 156), according to previous reports (20,46,47).

When accounting for clonal expansion and modelling Q1 condition (ii) (**Figure 1A**), the HLA effects on highly expanded clones were overtaken by association signals in HLA-DRB1 and DQ (**Figures 3E, S12E**). Strikingly, leading *cdr3*QTLs on expanded clones in healthy cohort overlapped with HLA loci associated with increased risk for CD (**Figure 3F, S12**) (19), shredding some light on the Q2 (**Figure 1A**) regarding functional consequences of IBD HLA risk loci. This convergence suggests that risk HLA alleles may influence IBD pathogenesis through *cdr3*QTLs, impacting composition of thymically selected and later expanded TRB clones.

### Universal landscape of HLA-DRB1 effects and disease-specific *cdr3*QTLs in HLA-B

Because of the strong linkage disequilibrium among polymorphic sites within the HLA regions, we applied a *cis*-conditional haplotype analysis using stepwise multivariate multiple regression, in which *cdr3*QTLs were sequentially added to the model to identify independent signals within a given HLA protein (27,37). Briefly, within each HLA protein, we started with the most significant *cdr3*QTL signal as an initial covariate during step-forward regression to the existing CDR3-QTL model and repeatedly tested multiple rounds of conditioning until we could not observe statistically significant signals within each of the HLA class I/II proteins (**Methods**).

Conducting conditional *cis*-haplotype analysis on combined cohorts, allowed us to reach the highest statistical power to capture HLA effects on CDR3 in both health and disease at the high resolution. We identified 54 novel independent *cdr3*QTLs: 14 sites in HLA-B and 40 sites in HLA class II, including 7 novel sites in HLA-DRB1 and 33 in HLA-DQ, all meeting a stringent significance threshold (P value < 3.7×10^⁻7^ = 0.05/136,982 total tests) (**Figures 4A, S14, S15**), to the best of our knowledge for the first time described here in the current study (27,30). Structurally, these sites were located in the peptide-binding pockets, such as HLA-B sites 114 and 156, both in E pocket, HLA-DRB1 site 67 in the P7 pocket, together with HLA-DQB1 sites 9 and 57, both in P9 pocket (**Supplementary Table 1**, **Figure S9**) (25). Both HLA-DRB1 site 67 and HLA-B 156 are surrounded by TCR contacting residues (**Figures S15, S16**) (48). In addition, polymorphisms in the HLA-DRB1 67–74 region contribute to TCR binding specificity (**Figure S16**) (49).

We also observed a CDR3β length-dependent *cdr3*QTL signals. The *cdr3*QTLs in the HLA-B had the strongest effects on CDR3 with lengths 12-14 amino acid long, which is shorter compared to the CDR3 length observed in HLA class II *cdr3*QTLs (**Figures S14A).** Leading *cdr3*QTLs, such as DRB1 67 and 71, HLA-B 156 and 116, and HLA-DQB1 sites 87 and 125, as well as HLA-DQA1 sites 129 and 156, had the strongest effect on the CDR3β composition, depending on the CDR3β length (**Figure 4, S15**). In addition to sites in the HLA-DQA1, leading *cdr3*QTL signals, predominantly, resided in peptide-binding pockets, suggesting *cdr3*QTLs influence the TRB repertoire via a peptide-presentation based route.

**Figure 4.**
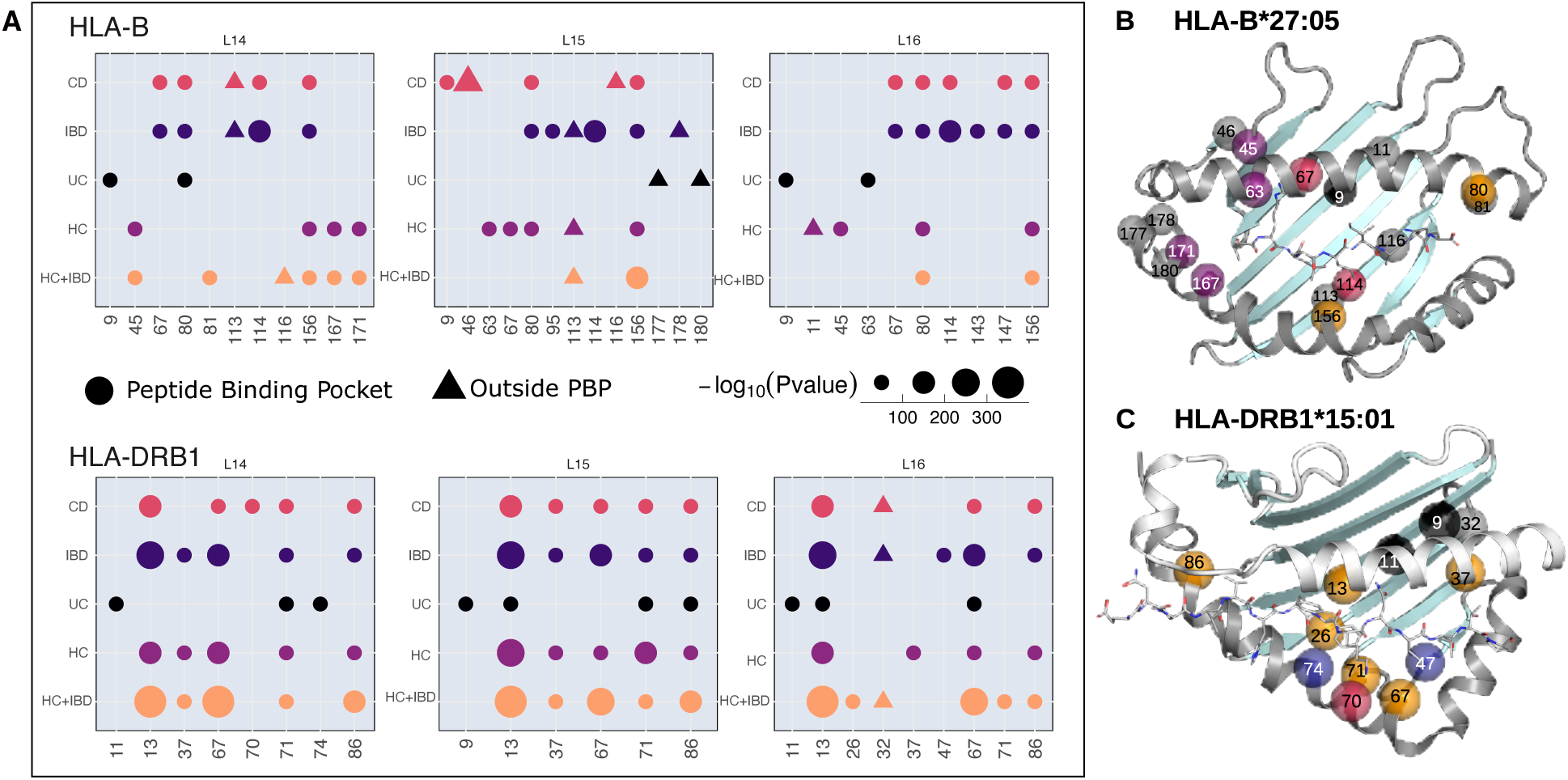
HLA fine-mapping of cdr3QTLs applying conditional cis-haplotype analysis and their localisation in HLA-peptide-TCR complexes. **(A)** Conditional haplotype analysis for healthy donors and individuals with IBD for HLA-B and HLA-DRB1, with the most common length being CDR3β 14-16. **(B)** Structural analysis of *cdr3*QTLs in HLA-B. Residues in the buried beta sheet are in contact with the peptide. Sites on the alpha-helices may contribute to TCR binding. **(C)** Structural analysis of *cdr3*QTLs in HLA-DRB1. Localisation of DRB1 cdr3QTLs in the protein structure of DR15/MBP peptide/TCR (PDB ID 1ymm). Most cdr3QTL sites are buried in the peptide binding groove, either deeply on the DRB1 central beta sheet (DRB1:13, 32, 37) or on the helix (DRB1:67, 71, 86). Some sites contact the peptide directly (N-terminal P1: 86; central P4-P7:13, 67, 71; C-terminal P9: 37). In DR15-TCR, no residues interact with the TCR, while in DR4-TCR sites 67, 70 and 71 do have contact with CDR3 110/111. Sites 67 and 71 are located at structurally flexible regions that adapt to the interacting TCR. Sites 13 and 86 are involved in structural changes during peptide processing. The specificity and efficiency of molecular dynamics are defined by polymorphisms at these sites. TCRα is in orange, and TCRβ is in yellow. Protein structural localisation of all independent *cdr3*QTLs in HLA-B*27 and HLA-DR15 peptide binding domains. Cα atoms of the sites are coloured as in (A). Most *cdr3*QTL sites are buried in the peptide binding groove, either deeply on the central beta or on the flanking helices. Only a few residues are not in direct contact with the peptide. HLA-B 156 and 171, and HLA-DR 67 and 86 co-locate in 3D, respectively. In HLA-B, alpha1 helical sites in HLA-B (63, 67) as well as 45 are in contact with the N-terminus of the peptide (pockets A and B), while in the beta sheet, sites are part of pockets B-F. In DRB1, *cdr3*QTL are contributing to peptide binding pockets P4, P6, and P9, and position 86 contributes to pocket P1. Some HLA-B sites may interact with TCR (63, 80, 167) or are neighbours of variant TCR contacting sites (67, 156, 171), while in HLA-DR, the alpha-helical kink region (position 67-74) is structurally flexible and sites 67, 70 and 71 are in contact with TCR in HLA-DR4 CDR3 110/111.

We aimed to explain the prominent CDR3β compositional differences between groups (**Figure 2B**), also reported in previous studies (8,50–52), by detecting disease-specific *cdr3*QTLs. After applying conditional *cis*-haplotype analyses on stratified cohorts, when statistically possible (**Figure S9**), we observed a clear segregation of *cdr3*QTLs between the IBD and non-IBD groups in HLA-B (**Figures 4**, **S15**). Specifically, HLA-B sites 9, 67, 113 and 114 were found exclusively in IBD patients, predominantly in CD patients, and sites 45 and 167 were found exclusively in the dataset of healthy controls. Nonetheless, HLA-B sites 80 and 156 had shared effects among stratified and combined groups. These HLA-B cdr3QTLs are located in deeply buried (9, 45, 114) and less buried (67,156, 167) peptide-binding pockets (**Figure 4B**). Of the latter, site 167 has a dual role by also selectively binding TCRs (53), and 67 and 156 are surrounded by TCR contacting residues (48). Since sites 9, and 156 shape *expanded singletons* CDR3β, antigen dependency may be suggested.

In contrast to segregated *cdr3*QTLs in HLA-B, six independent *cdr3*QTLs in HLA-DRB1 exhibited widespread “universal” effects independent of the disease, influencing the CDR3β values of 14 and 15 amino acids in both the IBD and non-IBD samples (**Figure 4A**). We validated the HLA-DRB1 *cdr3*QTL with permutational analysis of variance (PERMANOVA), which accounts for the compositional nature of both HLA and CDR3 (**Figure S19**). Given that *cdr3*QTLs in HLA class I and II molecules are involved in shaping “*naïve”*, or *expanded singletons* and expanded, hence antigen-experienced clones, with leading signals overlapping predominantly with GWAS CD risk loci (e.g., DRB1 sites 67, 71, 74) (20,39), this strongly implies that HLA molecules play a dynamic role in modulating T-cell responses implicated in IBD.

A structural analysis revealed that most of the *cdr3*QTLs in HLA-DRB1 and HLA-B exert their effects via peptide presentation, whereas some might have direct or indirect interactions with T cell receptors (**Figures 4, S16**). HLA-DRB1 sites 67-71 are found in a helical kink region that undergoes drastic structural rearrangements during HLA-DM-catalysed peptide processing and TCR binding (54,55). Further, HLA-II/peptide-TCR complexes may show unconventional topology as is the case for DR15/TCR with myelin basic protein (PDB IDs 1ymm)(56). Here, there is no TCR direct contact to *cdr3*QTL sites, but in the complex of DR4-enolin, the TCR is in contact with 67, 70 and 71, and this contact is established via the CDR3β positions 110 and 111 (57). Further, it was recently demonstrated that the deeply buried peptide-binding polymorphic sites also contribute to HLA structural dynamics in the process of DM catalysed peptide loading (58,59).

### HLA risk alleles associated with immune-mediated diseases cluster together based on their influence on the CDR3 composition

Building on the identified *cdr3*QTLs in HLA proteins, together with the fact that variations at some of those sites is genetically associated with several IMIDs, we hypothesised that HLA alleles with similar physicochemical properties at the *cdr3*QTLs site would form distinct clusters. Specifically, several DRB1 *cdr3*QTLs, *e.g.* 13, 67, 71, 74 have been linked with rheumatoid arthritis (RA), multiple sclerosis (MS), systemic lupus erythematosus (SLE), type 1 diabetes (T1D). Similarly, HLA-B sites 9, 45, 67 have been linked with RA, dermatomyositis, psoriasis and Grave’s disease (20,39,46). We found that all major HLA-DRB1 risk alleles for IMIDs clustered together on a single branch, due to very similar physiochemical properties (such as charge, size, and hydropathy) of their residues on key *cdr3*QTLs (**Figures 5, S20**). Within this unified “risk” branch, they are further partitioned into two distinct clusters (**Figure 5A**). Specifically, one cluster predominantly comprises HLA alleles strongly associated with type 1 diabetes (T1D) and rheumatoid arthritis (RA), such as DRB1*04:01/04 (20). The other cluster contains other strong risk alleles, such as DRB1*15:01, associated with multiple sclerosis (MS), systemic lupus erythematosus (SLE), and DRB1*01:03, associated with risk for both the IBD subtypes CD and UC (19), all of which share common underlying physiochemical characteristics (**Figure 5B**). Intriguingly, DRB1*03:01, a prominent risk allele for SLE, T1D, and Graves’ disease, clustered unexpectedly with DRB1*13:01, a protective allele for T1D, MS, and RA, but is a risk factor for the autoimmune liver disease, primary sclerosis cholangitis (60). This physiochemical proximity of residues at *cdr3*QTL might suggest unique functional implications and distinct underlying structural mechanisms that differentiate risk and protective HLA alleles.

The role of HLA-B alleles in IMIDs is less prominent than that of some HLA class II alleles. Nonetheless, we observed that signals within HLA-B were associated with the CDR3 composition of *expanded singletons*. HLA-B risk alleles for ankylosing spondylitis (HLA-B*27:05) (20,61), and type 1 diabetes (HLA-B*39:01/06 (20,62), HLA-B*18:01 and HLA-B*13:01/02 (20)) clustered tightly together, showing distinct effects on CDR3 linked to those alleles (**Figure S20A-B**). We inferred CDR3 motifs from public TRB clones that were significantly linked to HLA-B alleles within each cluster (**Figure S20C-E**). Compositions of CDR3 clones linked to one cluster were dissimilar from those, linked to another cluster (**Figures 5D, S20**). These motifs highlighted recurrent TCR patterns and HLA effects on the CDR3 loops, potentially mediated by antigens, as key *cdr3*QTLs localise to peptide-binding pockets.

**Figure 5.**
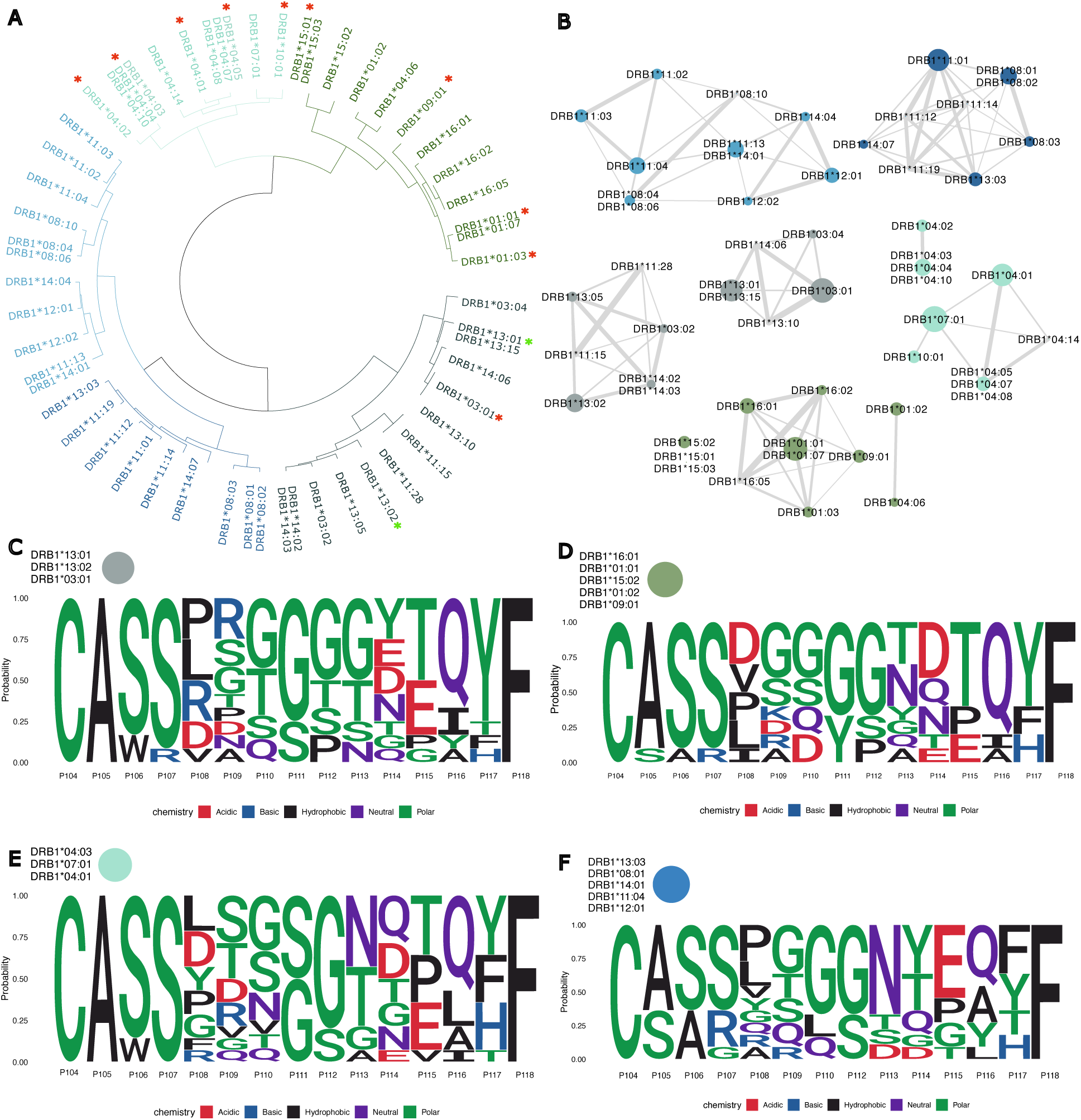
Clustering of HLA-DRB1 alleles, based on the physicochemical properties of amino acids, found at key cdr3QTLs with subsequent network analysis. **(A)** Common HLA alleles from the European population formed five distinct clusters (colour-coded) from two main branches. **(B)** Network analysis, which is based on amino acid physicochemical similarities at cdr3QTLs, revealed that the key disease risk-susceptible HLA alleles for MS, SLE, RA, T1D, UC and CD clustered on one branch, with separation of clusters between MS, SLE, UC, and CD in one cluster and T1D with RA in another cluster. The key protective alleles of the DRB1*13 family and a risk allele for T1D, RA and SLE DRB1*03:01 share similar amino acid properties at *cdr3*QTLs and are located on another branch. **(C)** CDR3 motifs of public TRB linked to protective HLA alleles and those associated with CD. **(D)** CDR3 motifs of public TRB linked to HLA alleles associated with RA and MS. **(E)** CDR3 motifs of public TRB linked to HLA alleles associated with RA and T1D. **(F)** CDR3 motifs of public TRB linked to protective HLA alleles and those associated with RA.

## Discussion

The increased global prevalence of IBD underscores the critical need for a deeper understanding of its complex aetiology, particularly the interplay between genetic predispositions and dysregulated immune responses. While the functional consequences of HLA variations in autoimmune and chronic inflammatory diseases have been difficult to identify (18,19), emerging evidence suggests their crucial role in shaping T-cell receptors (26,27,30,43). Earlier studies highlighted the selective role of some HLA alleles and variable sites on TCR repertoires through biased usage of germline-encoded V and J genes (26,43) and the selection of specific chemical properties, lengths and composition of CDR3 regions (27,28,43).

By conducting a CDR3-QTL analysis with multiple expansion-weighted strategies across >1,900 samples from healthy individuals and those with IBD, we explored the effects of HLA on TCRs during T-cell activation in the periphery and positive and negative selection in the thymus. We observed that HLA-DRB1 exerts effects on the CDR3 repertoires of *singletons* in both healthy and IBD-affected individuals through the same set of *cdr3*QTLs with effects stronger than the phenotype effects (**Figures 2D, 4A**), suggesting the existence of HLA-DRB1 universal influences on the composition of TCRs irrespective of the disease. On the other hand, *cdr3*QTLs in HLA-B play a more dynamic, IBD-dependent role in shaping the amino acid composition of the CDR3s (**Figures 3C, S12**). IBD-specific *cdr3*QTLs in HLA-B (**Figure 4A**) were associated with distinct shifts in CDR3β characteristics exclusively within the IBD cohort. Interestingly, IBD-specific *cdr3*QTLs in HLA-B overlapped with sites that modulate responses to viral infections *in vivo* (46,47), for example, HLA-B sites 9 to EBV and VZV and sites 67 and 156 to HIV. This may explain the increased risk of IBD after HIV infection (63), which leads to CD4^+^ T-cell depletion and the involvement of CD8^+^ T cells in IBD pathogenesis. Murine *in vivo* models have shown that cytotoxic CD8^+^ T cells induce relapsing colitis with subsequent remission after anti-CD8 therapy (64,65). The effects of HLA-B on TRB *singletons* offer insights into the earliest stages of immune dysregulation (8,52) and disease susceptibility rather than merely reflecting the consequences of chronic inflammation.

In contrast to the abovementioned effects, *cdr3*QTLs in HLA class II have minimal effects on the CDR3 composition of the most expanded clones when we focus on their composition without considering their expansion, referred to as *expanded singletons*. This is remarkable, as HLA class II had a strong effect on the composition of the top expanded TRB clones when accounting for clonal expansion. The most significant HLA-DRB1 and HLA-DQA1 effects on highly expanded TRB clones and *expanded singletons* within healthy controls, and not within IBD, overlapped with the main GWAS risk loci for CD (**Figure 3E**), whereas the leading *cdr3*QTLs, associated with the CDR3 composition of singletons of both the IBD and HC repertoires, reflecting HLA effects that occur in the earliest stages of T-cell development, overlap with both CD and UC risk loci. This highlights different mechanisms of HLA genetic risk for CD and the dynamic interplay between predispositions in healthy individuals, HLA-driven thymic selection of T cells, clonal expansion, and potential disease onset. In line with this, a recent study has shown that the nature of infection, whether acute, latent, or chronic, modulates T-cell phenotypes, even when the same antigenic epitope is targeted (66). Taken together, our results highlight that both genetic and environmental factors shape disease-relevant immune repertoires.

Our results suggest the existence of different mechanisms in health and disease, together with some universal, irrespective of disease, principles of HLA-driven selection of T-cell receptors. Given the location of many *cdr3*QTLs within peptide-binding pockets, our data suggest a peptide-mediated model of the effects of HLAs on the immune repertoire (**Figures 4, S16**). We could not separate the independent contributions of *cdr3*QTLs to each of the maternal and paternal HLA haplotypes or the additive effects of their interactions, due to the lack of phasing information.

Finally, we observed that key HLA risk alleles associated with multiple inflammatory diseases, including RA, T1D, SLE, MS, and UC, but not CD, clustered together on the basis of similarities in key *cdr3*QTL physicochemical properties, leaving the main protective alleles in the opposite branches. This suggests a long-standing evolution behind the vast diversity and functional consequences of HLA biology (**Figure S20**) (67,68). The observed selection pressure on HLA-B alleles encoding an isoleucine at position 80 in response to *Y. pestis* infection in ancient human populations (69) is particularly compelling, as HLA-B site 80 was among the identified *cdr3*QTLs, strongly influencing the amino acid composition of the T-cell receptor (TCR) repertoire. This opens the question about the evolutionary success of certain HLA alleles in response to common infections: was it a direct selection for the HLA molecule itself, or rather a byproduct of selecting for the generation of a T-cell repertoire with CDR3s optimally suited to combat the pathogens?

## Conclusion

Our study revealed that specific sites in HLA class-I/II proteins, *i.e. cdr3*QTLs, influence the composition of the TCR receptors utilised by αβ T cells. Given the evidence that HLA sites are highly effective in shaping T-cell repertoires against common environmental (47,70,71) and/or self-antigens, we propose that *cdr3*QTLs act as key determinants in the positive selection and may, under particular circumstances, such as environmental exposures, drive IBD pathogenesis through their effects on the amino acid composition of T-cell receptors. This fine-grained influence has not been previously shown, especially in the context of chronic inflammatory diseases, such as IBD. The tight haplotype structure between HLA genes is a well-known problem in genome-wide association and fine-mapping studies, and pinpointing the causal and independent HLA genetic effects is still a major challenge (21,72). Future studies on paired αβ T-cell repertoires with phased genetic architectures investigating the precise impact of HLA-driven variations on T cells, the antigenic specificities of these T cells, and the molecular mechanisms of complex interplay between host genetics, heterogeneous T-cell populations, and environmental triggers is highly needed.

## Abbreviations

CDR3: complementarity-determining region 3
TRB: T-cell receptor beta chain
TCR-seq: T-cell receptor sequencing
HLA: human leukocyte antigen
IBD: inflammatory bowel disease
CD: Crohn’s disease
UC: ulcerative colitis
HC/HBD: healthy controls/healthy blood donors
RA: rheumatoid arthritis
MS: multiple sclerosis
PBS: peptide binding site (HLA)
PSC: primary sclerosing cholangitis

## Acknowledgements

The project was funded by the EU program for Research and Innovation “Horizon Health” (HORIZON-HLTH-2023-DISEASE-03) ID-DarkMatter-NCD (897856542). Additionally, the project received funding from the German Research Foundation (DFG) Research Unit 5042: miTarget – The Microbiome as a Therapeutic Target in Inflammatory Bowel Diseases, along with funding from the DFG Cluster of Excellence 2167 “Precision Medicine in Chronic Inflammation (PMI)”.

