## Supplemental Appendix for "The dynamic role of HLA proteins on compositional alterations of T-cell repertoires in inflammatory bowel disease"

### Supporting Information Appendix

#### Extended Results

##### Conditional *trans*-haplotype analysis

In an attempt to resolve the independence of cdr3QTLs within the HLA class I and II regions, we performed conditional *trans*-haplotype analyses. Conditioning HLA-DQ signals on the basis of known DRB1-DQA1-DQB1 haplotypes would not provide any additional information to CDR3-QTL models because of the determined genetic nature of known haplotypes. As we did not have phasing information about the exact combination of HLA variants within an individual's chromosomes, we assessed pair-wise correlations of allele dosages between all HLA site allelic variants (**Figures S17, S18**) and used individuals' HLA site variant correlations ($r^2 > 0.6$ and p value $< 0.05$) as observed haplotypes. We observed the strongest correlations between HLA class II sites. Conditioning HLA-DQA1 sites on the strongest hits in HLA-DRB1: sites 13, 37 and 71 led to the model performance dropping to the negative values, meaning that three HLA-DRB1 sites explained CDR3 composition better than with any additional HLA-DQA1 site.

### Supplemental Figures

### S1: Distribution of HLA alleles across healthy blood donors and IBD patients

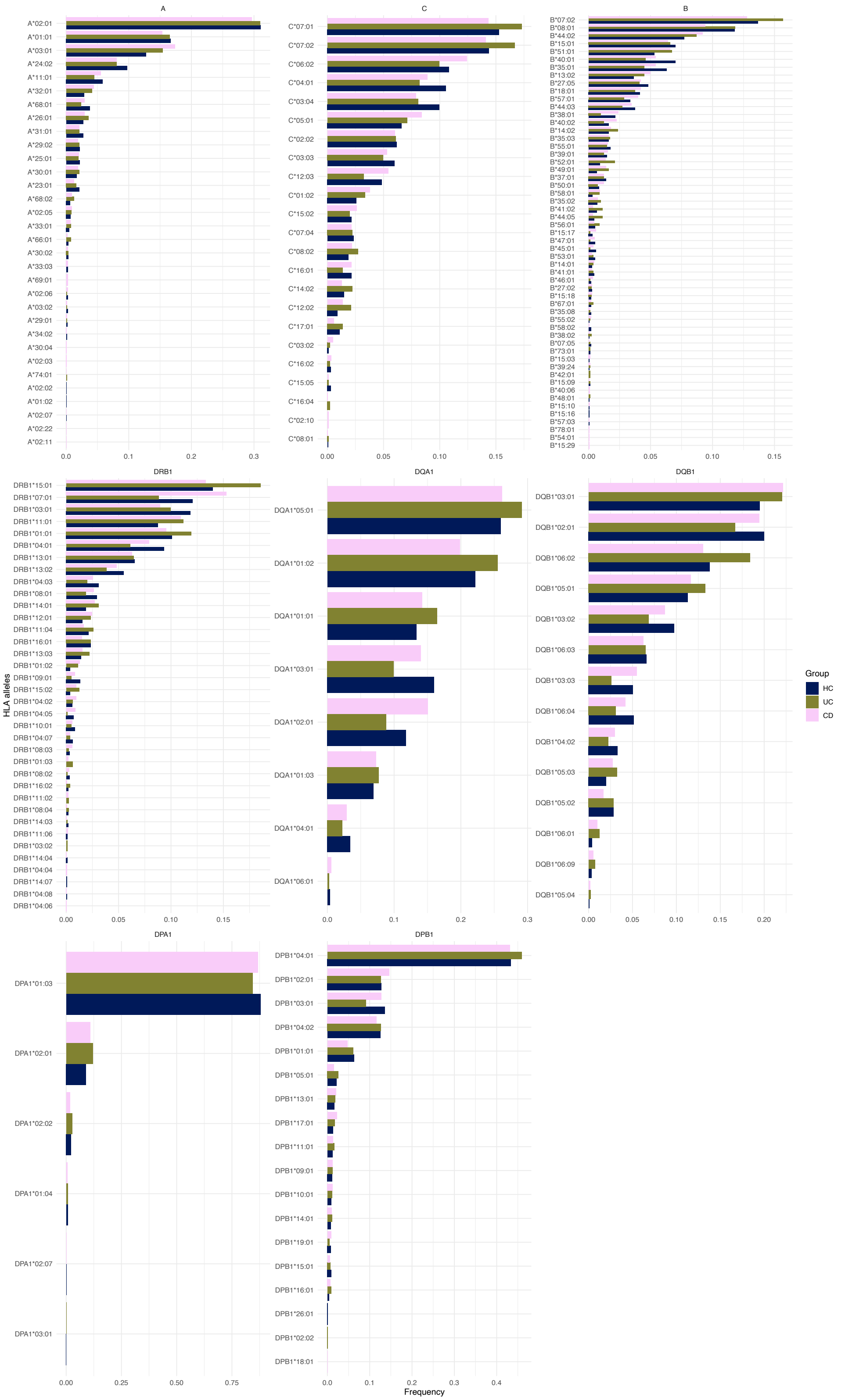


**Figure S1**. HLA class I and class II alleles frequencies across cohorts. Bars represent the observed frequencies (on X axis) of specific HLA alleles (Y axis) within each study group.

#### S2: The TRB repertoire analysis

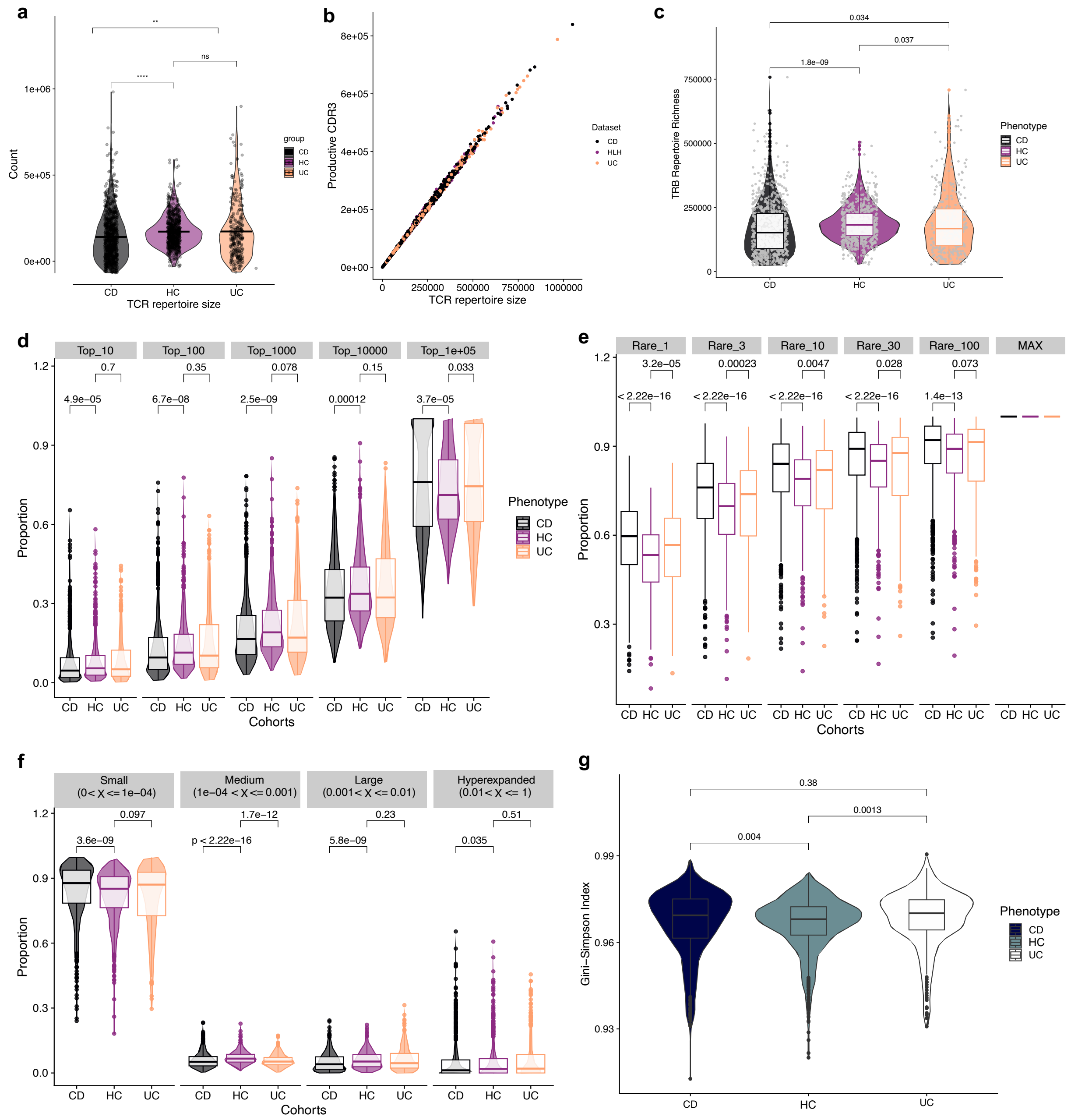


**Figure S2.** *The TRB repertoire analysis.* A: Comparison of repertoire sizes across studies groups. B: The proportion of productive receptor sequences from the bulk TCR-seq data. C-F: Richness, diversity and clonality of immune repertoires provide a robust overview of T cell antigenic properties in healthy states and diseases (47–50). C: Richness of repertoires. D: Clonality across top 10, 100, 1000, 10e3, 10e4 expanded clones, on Y axis is the proportion that expanded clones occupy from the whole repertoire, and phenotypes on the X axis. E: Low-to-moderate clonal proportion of rare clones, metric that captures the part of the repertoire made up of clones with relatively low expansion levels (from singletons up to 100 copies). F: Repertoire clonal homeostasis: it displays the proportion of rare, medium, large and hyperexpanded clones within the repertoire. G: Gini-Simpson index as a metric of evenness and diversity.

**S3: Distribution of public CDR3s**

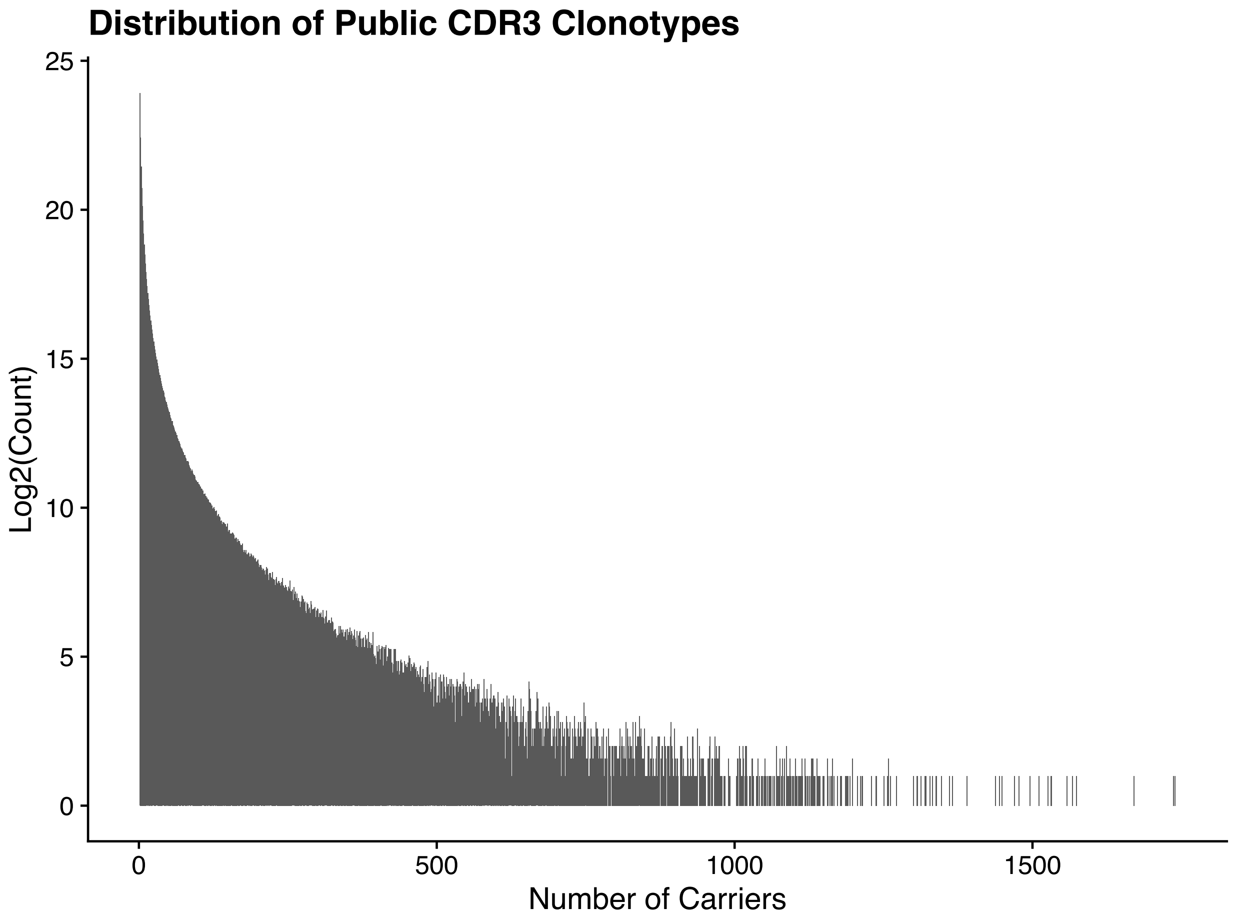


**Figure S3.** Each bar represents a unique public CDR3 clonotype. The X-axis indicates the number of individuals sharing that clonotype (number of carriers). The Y-axis represents the log2-transformed total abundance of each clonotype across the entire cohort, illustrating that unique clonotypes tend to exhibit higher overall abundance.

#### S4: CDR3 motifs of public clones, linked to specific HLA-B and HLA-DRB1 alleles

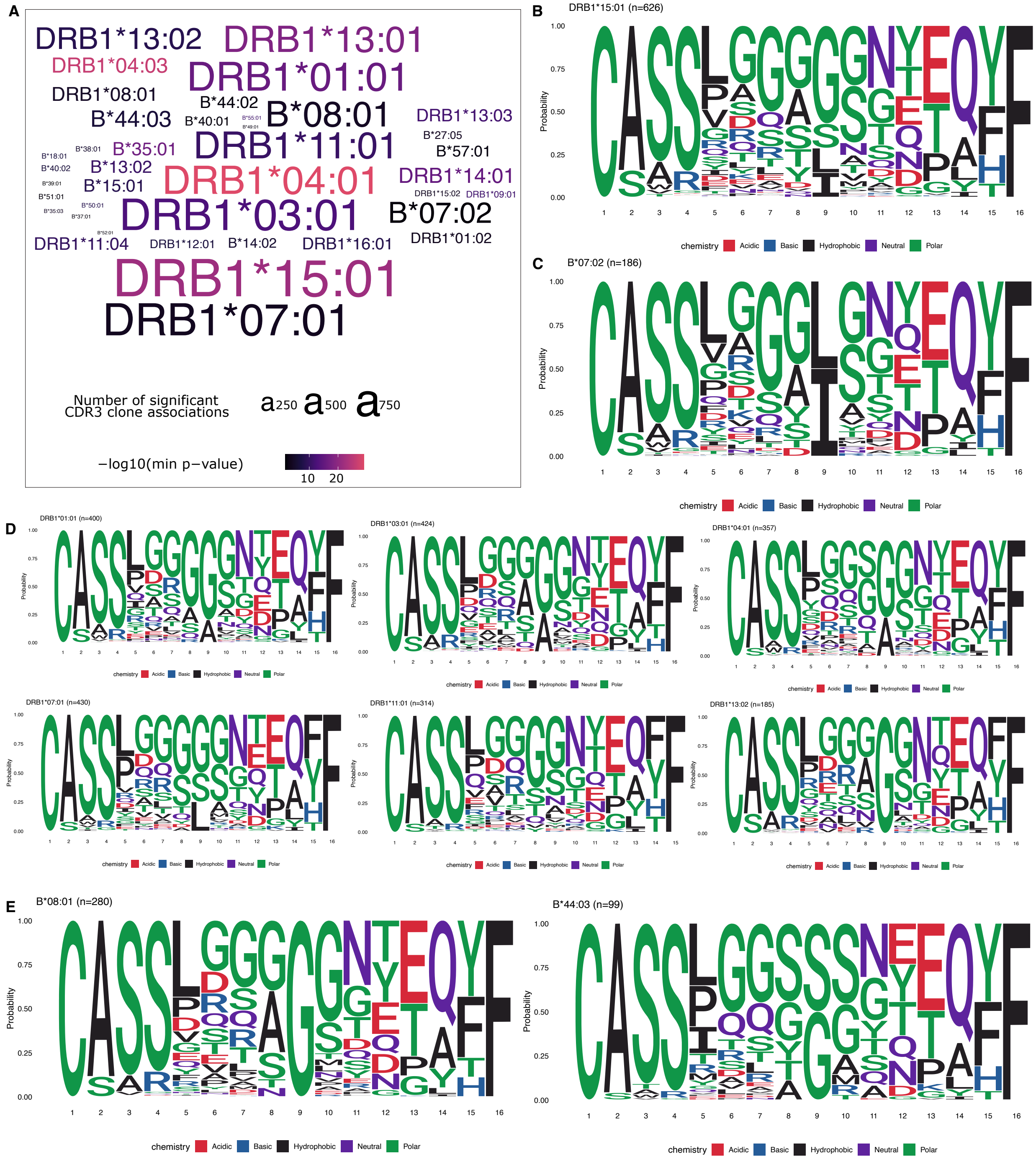


**Figure S4.** *HLA alleles leave detectable signatures on TRB public clones.* **A**: HLA-B and HLA-DRB1 alleles, associated with TRB clones, the font size stay for the number of associated clones, and colour shows the p value of the strongest association with certain HLA allele. B: Sequence logos illustrating the consensus amino acid motifs of public CDR3β clonotypes that demonstrated a significant statistical association with HLA-DRB1*15:01 allele. Each logo represents the amino acid preferences (measured in probability, Y-axis) at each IMGT position within the CDR3 region (1-16 represent the IMGT positions, starting from P104 to P118). Panel A-E: CDR3 motifs identified for public clonotypes significantly linked to common HLA-DRB1 alleles. These motifs highlight recurrent TCR recognition patterns and potential shared structural features within the CDR3 loops that mediate antigen presentation via specific MHC Class I and Class II molecules. Clonotypes included in motif generation met an adjusted P-value after Bonferroni correction threshold of $p < 0.05$ from the association analysis.

#### S5: The CDR3 amino acid composition differences

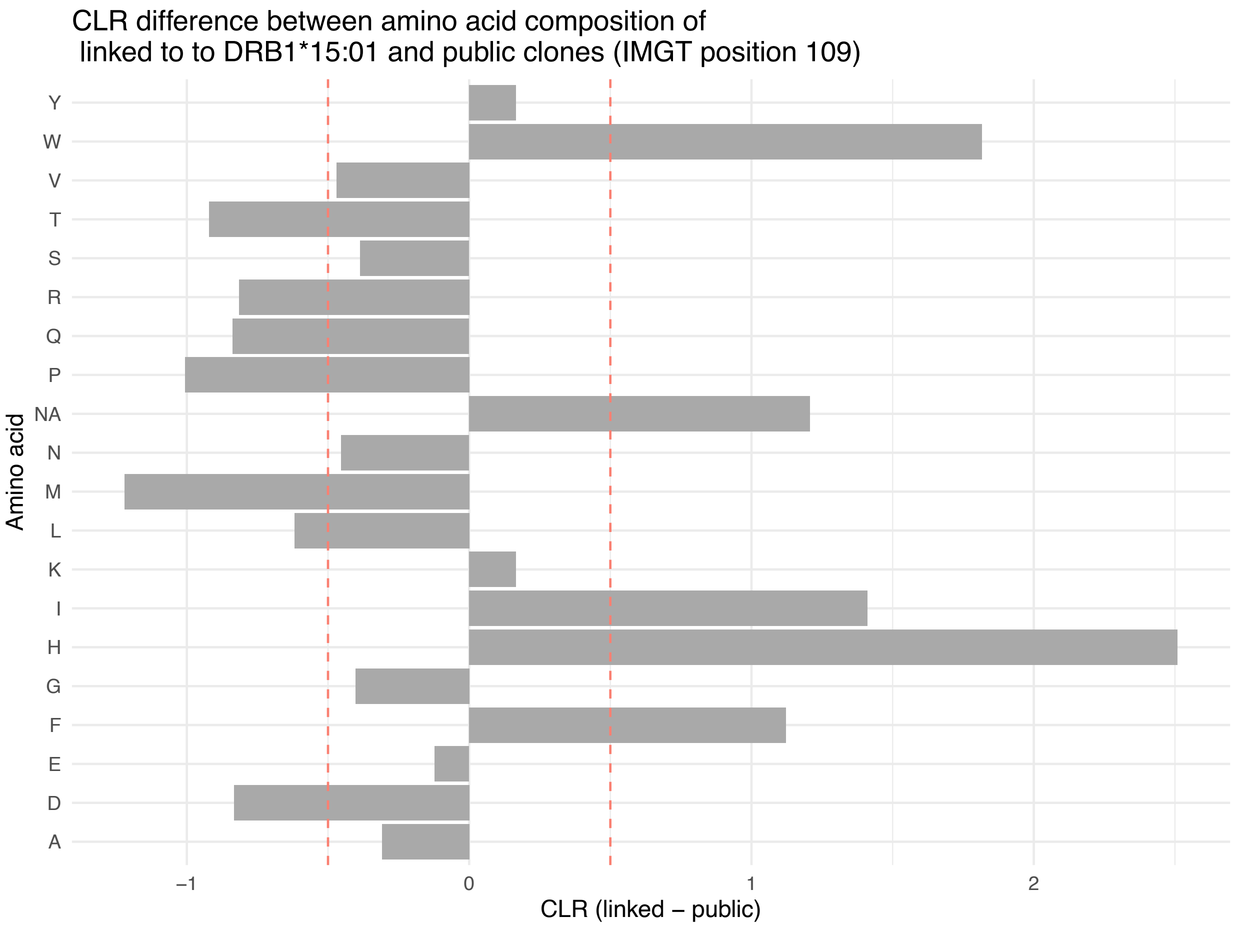


**Figure S5.** *Aitchison distance between amino acid compositions of public TRB clones and clones linked with HLA-DRB1*15:01.* Centered log-ratio (CLR) differences in amino acid composition at IMGT position 109 between TRB clones associated with HLA-DRB1*15:01 and public TRB clones. Positive values indicate amino acids that are more frequent in DRB1*15:01–linked clones, while negative values indicate amino acids that are more frequent in public clones. Red dashed lines mark the zero baseline (no difference).

#### S6: CDR3 motifs of public clones, linked to specific HLA-B allele clusters

#### S7: Site-specific HLA-DRB1 effects on CDR3

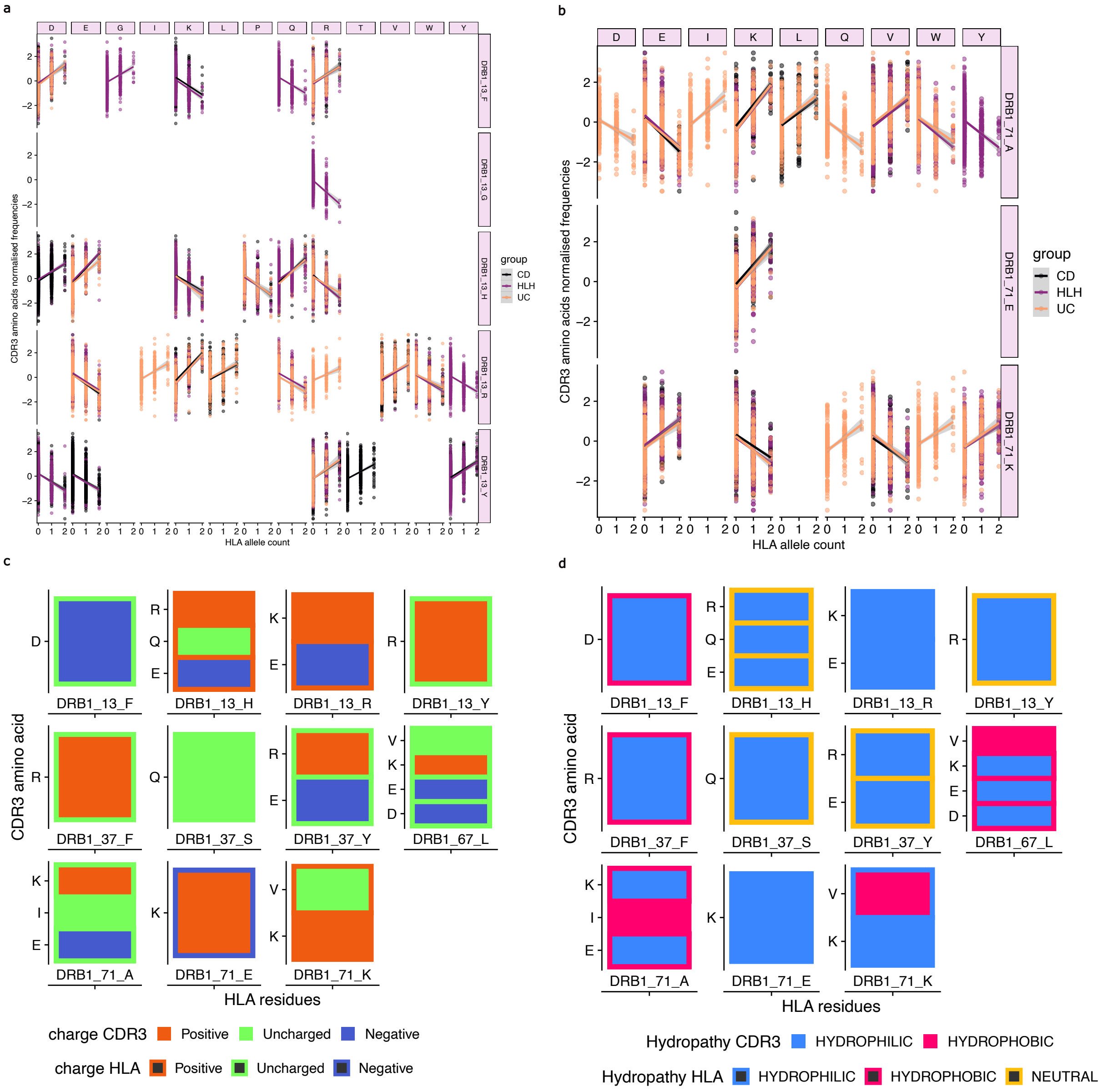


**Figure S7.** *Initial characterisations of HLA-CDR3 relationships on the example of DRB1 sites 13 and 71, previously shown to influence CDR3 composition.* A-B: We measured Pearson correlations between HLA allelic variant dosages at each variable HLA site and CDR3 amino acid frequencies at each position, according to IMGT nomenclature. C-D: Our analyses revealed a strong effect for hydrophilic and polar amino acids in CDR3β positions 110 – 112 with sequence lengths 14 – 17 amino acids long, consistent with the previous studies (2,3).

#### S8: Pearson correlation between HLA site variants and affected CDR3 amino acids

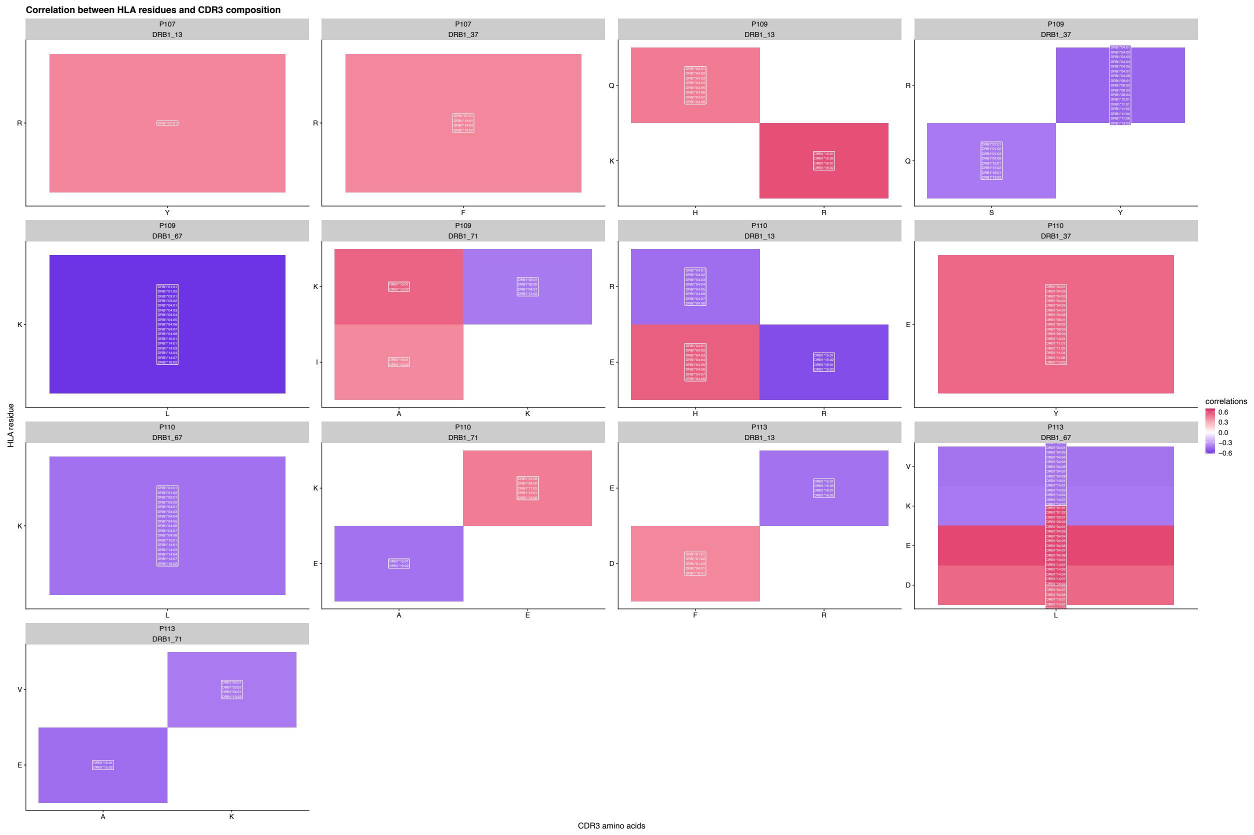


**Figure S8.** *Correlations between specific HLA site variants and the amino acid composition at their associated CDR3.* On the Y-axis the HLA amino acid (residues), while the X-axis correspond to specific amino acids within each non-template CDR3 position (P107-P113). The color gradient reflects the Pearson correlation coefficient for each pair (visualised only pairs with $r > |0.3|$ and p value $< 0.05$), with red indicating a strong positive correlation and violet indicating a strong negative correlation. This figure highlights the precise associations between key cdr3QTL HLA site variants residues and the resulting amino acid preferences in the T-cell receptor CDR3 loop.

#### S9: Serial downsampling

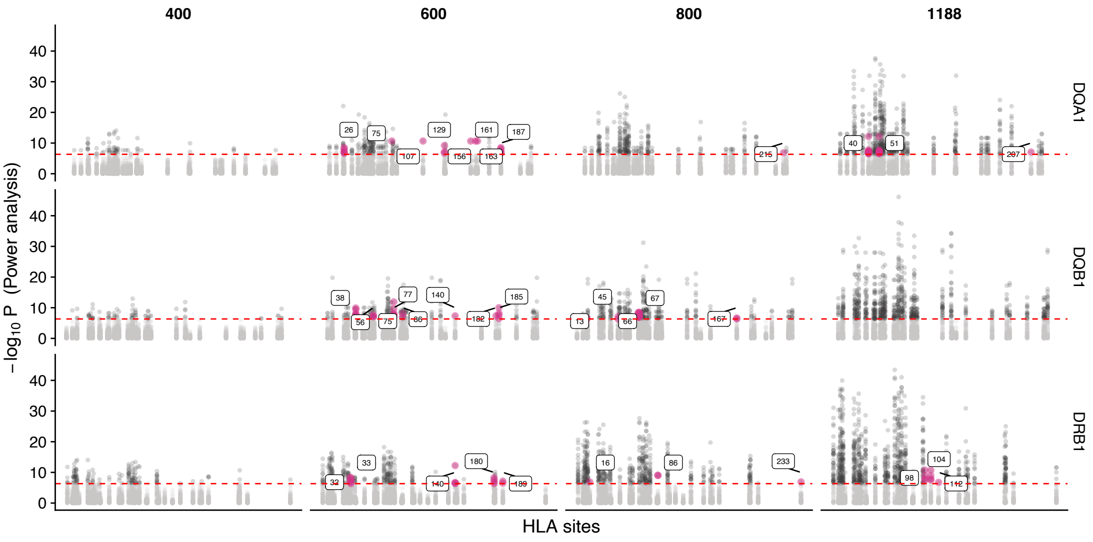


**Figure S9**. *Power Analysis for CDR3-QTL Discovery mapping in the IBD Cohort.* A: The relationship between increasing sample size and the detection of novel cdr3QTLs (depicted in pink). Results were derived from a power analysis based on the serial down-sampling of full IBD cohort (n=1188: 400, 600, 800, 1188). The X-axis represents the tested HLA sites, while the Y-axis shows the statistical significance of HLA-CDR3 associations, when CDR3 amino acid frequencies per each IMGT position were recalculated for each sample sample. The novel pink dots demonstrates that larger sample sizes are essential for enhancing statistical power and identifying a greater number of novel cdr3QTL associations.

#### S10: Comparing results between Healthy cohorts from current study to the earlier CDR3-QTL mapping

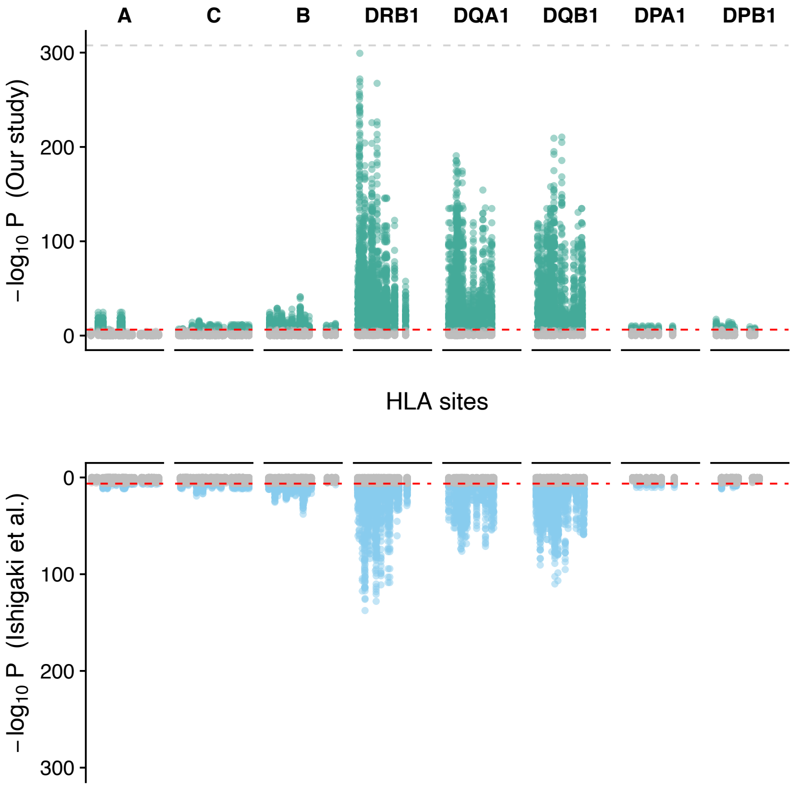


**Figure S10.** Increased power to detect CDR3-QTL associations with larger sample size for healthy cohort. This Miami plot compares the statistical significance of shared CDR3-QTL associations between the current study (N=772, Y-axis upper part) and the previously published study *(4)* (N=666, Y-axis lower part). Each point represents a CDR3-QTL association identified in both cohorts, with significance depicted as -log10 P-values. Our increased sample size provides substantially greater statistical power, leading to stronger detection of previously detected as well as novel *cdr3*QTLs (**Figure 2**).

#### S11: HLA sites at which phenotypes have a higher or lower explanatory power

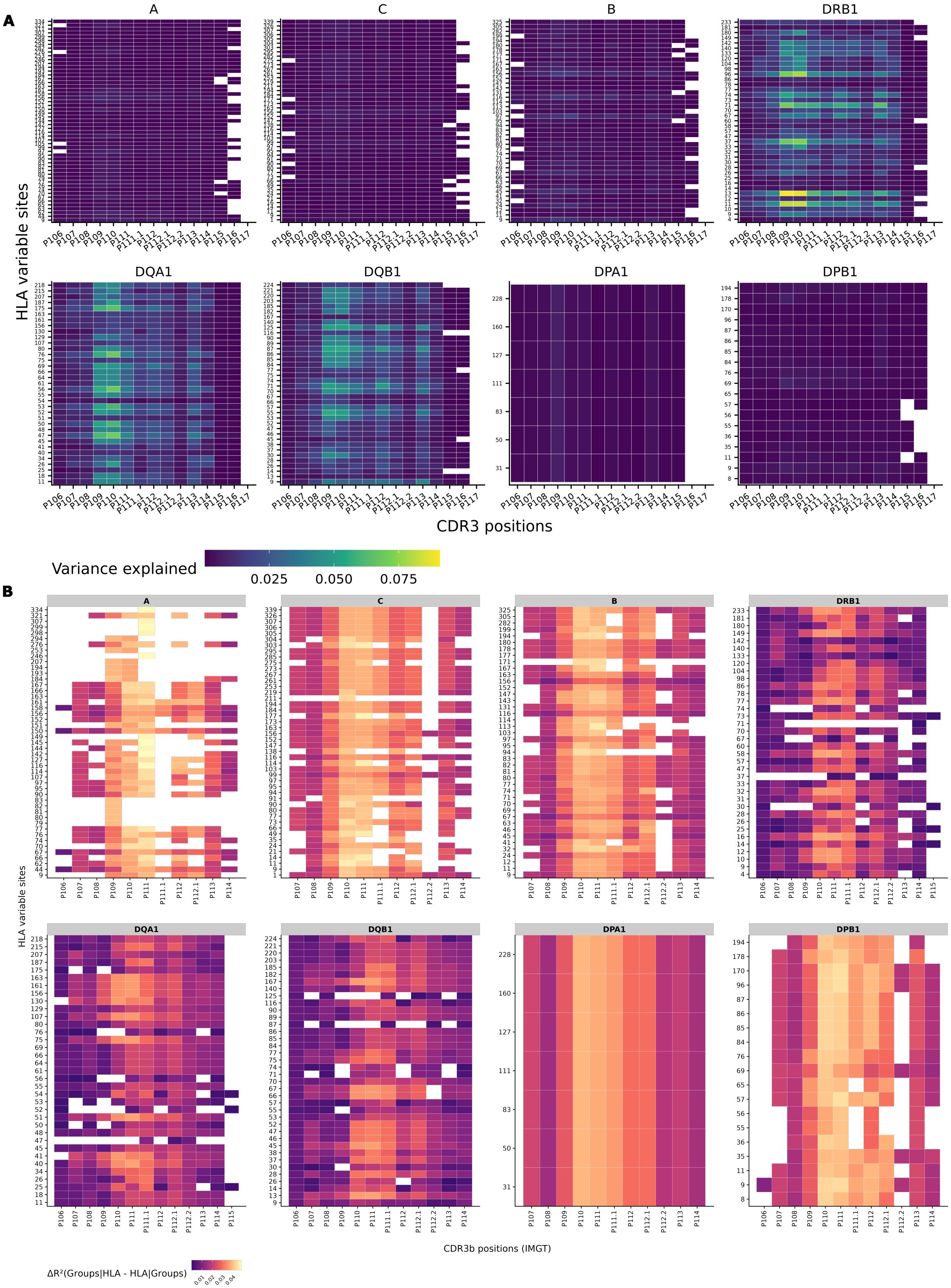

**Figure S11.** *The proportion of CDR3 amino acid composition variance explained by HLA sites and group phenotype ($R^2$).* **A:** This heatmap visualises the proportion of variance in CDR3 composition at the level of amino acid for each IMGT position (X-axis) explained by all HLA site variants matrix (Y-axis). The color intensity within each cell represents the proportion of variance explained for that particular association, with a gradient from blue (low variance explained) to yellow (high variance explained). This figure highlights the key HLA residues that significantly contribute to shaping the overall amino acid composition and diversity of the CDR3 repertoire. **B:** Heatmap showing the relative explanatory power of Groups conditional on HLA (Groups|HLA) across HLA sites (y-axis) and CDR3 positions (x-axis). Each tile represents the variance in CDR3 composition explained at the corresponding HLA–CDR3 site pair, with higher values indicating stronger group-specific effects after accounting for HLA.

#### S12: Distinct HLA-CDR3 associations in naive-like vs. antigen-experienced TRB repertoires and overlap with Crohn's Disease risk loci

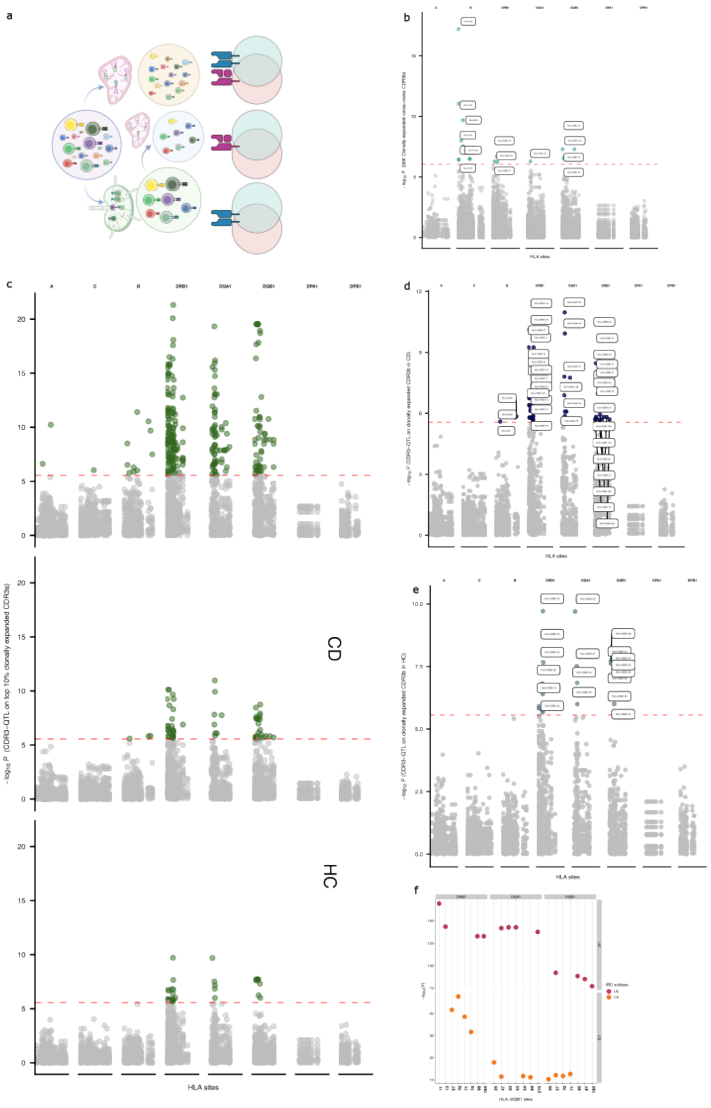


**Figure S12.** A series of TRB repertoire downsampling experiments with fixed numbers of clones ($3{\times}10^4$, $6{\times}10^4$, $10^5$, $2{\times}10^5$, $3{\times}10^5$) coupled with association testing between HLA site variants and CDR3 composition. (**A**): To differentiate between naive-like and antigen-experienced TRB clonotypes, the repertoire was analyzed under three conditions: "flattened to singletons" (putative naive, in yellow): by collapsing all clonotypes to a single count per sample, thereby emphasizing the HLA effects on unexpanded clonotypes. "Expanded" (putative antigen-experienced, in green circle): by focusing on highly expanded clonotypes and selecting a subset of reads that enrich for expanded clones. “Expanded-once naive” clones are those highly expanded, but after collapsing their expanded frequencies, focusing on their composition, but at earlier stages on clonotype development. (**B**): HLA *cdr3*QTLs (*e.g.*, significant associations between HLA sites and CDR3 compositional features) identified primarily in the downsampled to 30K “expanded singletons” repertoire. C: HLA cdr3QTLs identified in the "expanded" repertoire, segregated by cohorts. D: Annotated cdr3QTLs in HLA proteins found in expanded, antigen-experienced TRBs from IBD cohort. E: Annotated cdr3QTLs in HLA proteins found in expanded, antigen-experienced TRBs from healthy cohort. Critically, HLA cdr3QTLs discovered in the expanded, antigen-experienced TRB repertoire from healthy individuals demonstrated significant overlap with established HLA risk loci for Crohn's Disease (CD) (as indicated by (**F**). This finding suggests that specific HLA alleles shape the repertoire of antigen-experienced T-cells in ways that may predispose to or protect from inflammatory bowel disease. All associations with P-value < 0.05/26,880 (total number of tests) are colored in blue, green or purple, depending on the analysis, otherwise those not meeting the threshold were colored gray.

**S13: CDR3-QTL on expansion-weighted serially downsampled TRB repertoires of combined cohorts**

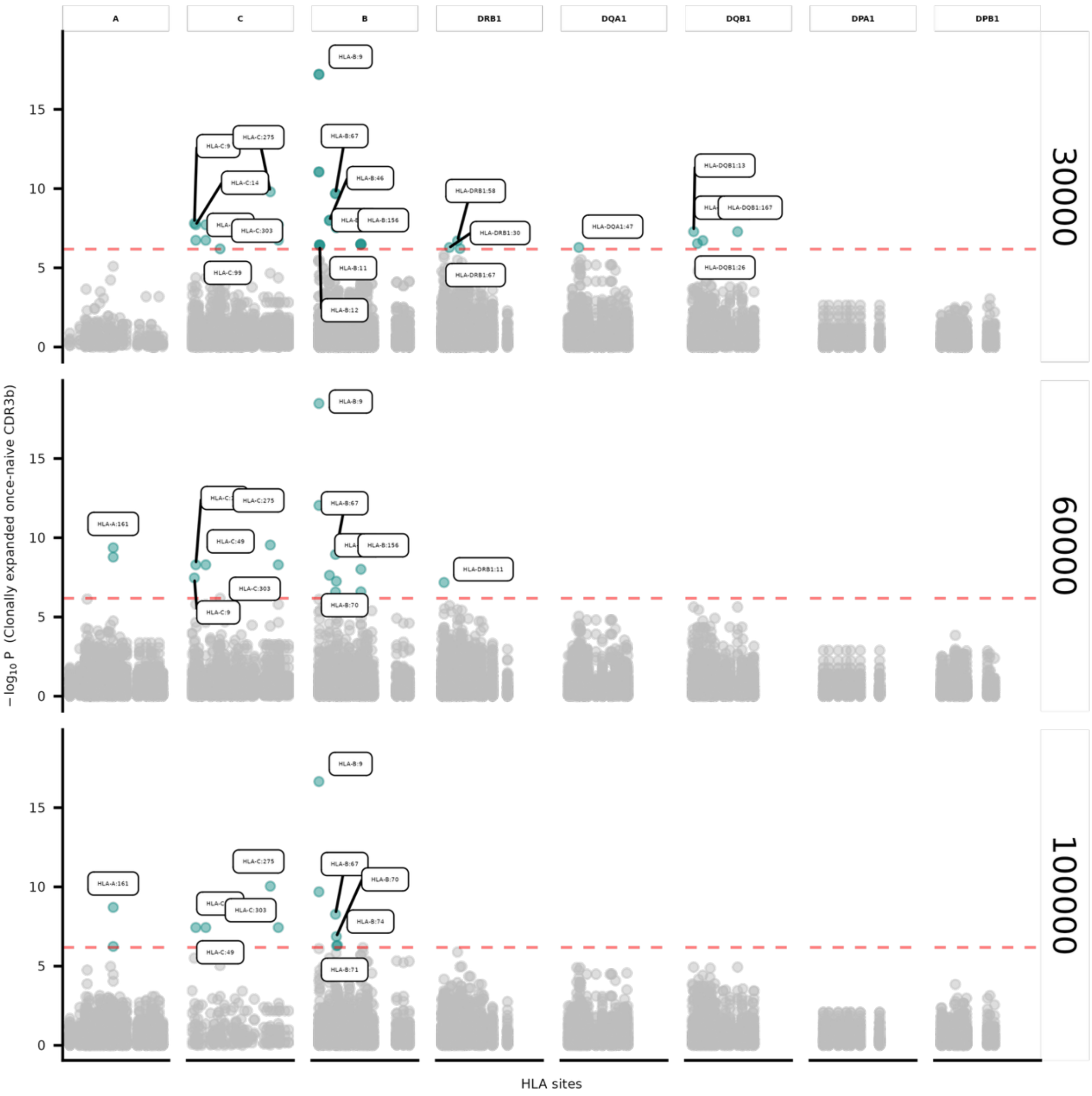


**Figure S13.** Serial down-sampling strategy of *expanded singletons*, based on increased probability of inclusion of a clone to be in the repertoire depending on clones' expansion. HLA-B *cdr3*QTLs shape initial amino acid composition of highly expanded clones.

#### S14: Conditional cis-haplotype analyses after CDR3-QTL on Combined IBD and HC singleton repertoires

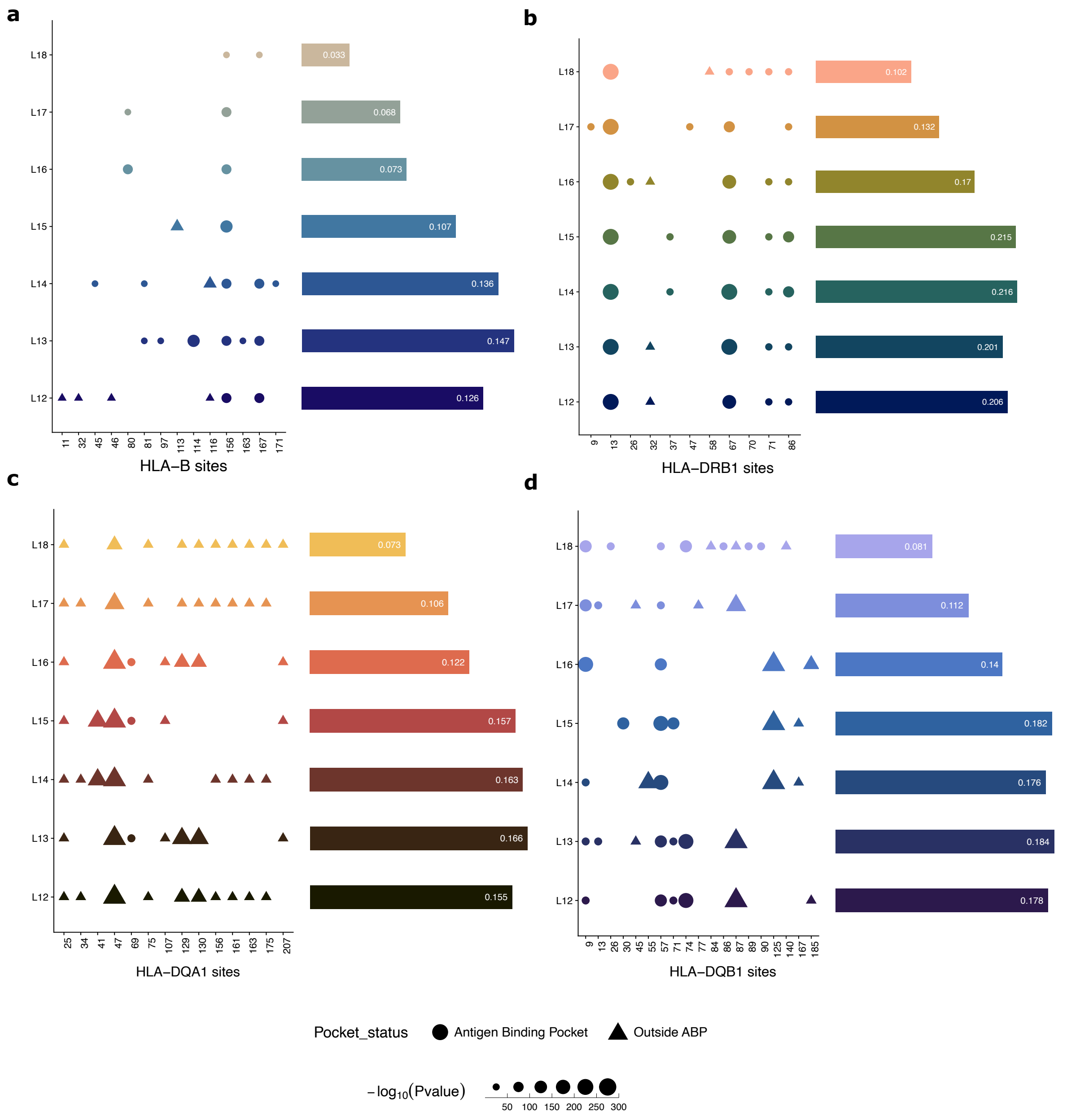


**Figure S14.** *Independent cdr3QTLs revealed by conditional cis-haplotype analysis on the combined dataset*. Each plot displays multiple HLA sites – cdr3QTLs, on the x-axis, corresponding to amino acid positions within the respective HLA chains. The y-axis indicates CDR3β lengths from 12 to 18 amino acids. Bars indicate the proportion of variance within CDR3 explained by particular combination of cdr3QTLss, with circles representing location in antigen-binding pockets (ABP), and triangles for those HLA sites outside of antigen-binding pockets. Sizes of shapes represent the strength of cdr3QTL associations (p-values). The figure reveals novel cdr3QTL signals within A: HLA-B cdr3QTLs, most of them reside in ABP; B: HLA-DRB1, which collectively explain over 20% of the variation in amino acid frequencies for CDR3β length 14-15; C: HLA-DQA1, and D: HLA-DQB1. A notable finding is the presence multiple cdr3QTLs within each DRB1 and DQ chains being shared across different CDR3 length.

#### S15: HLA map with *cdr3*QTLs

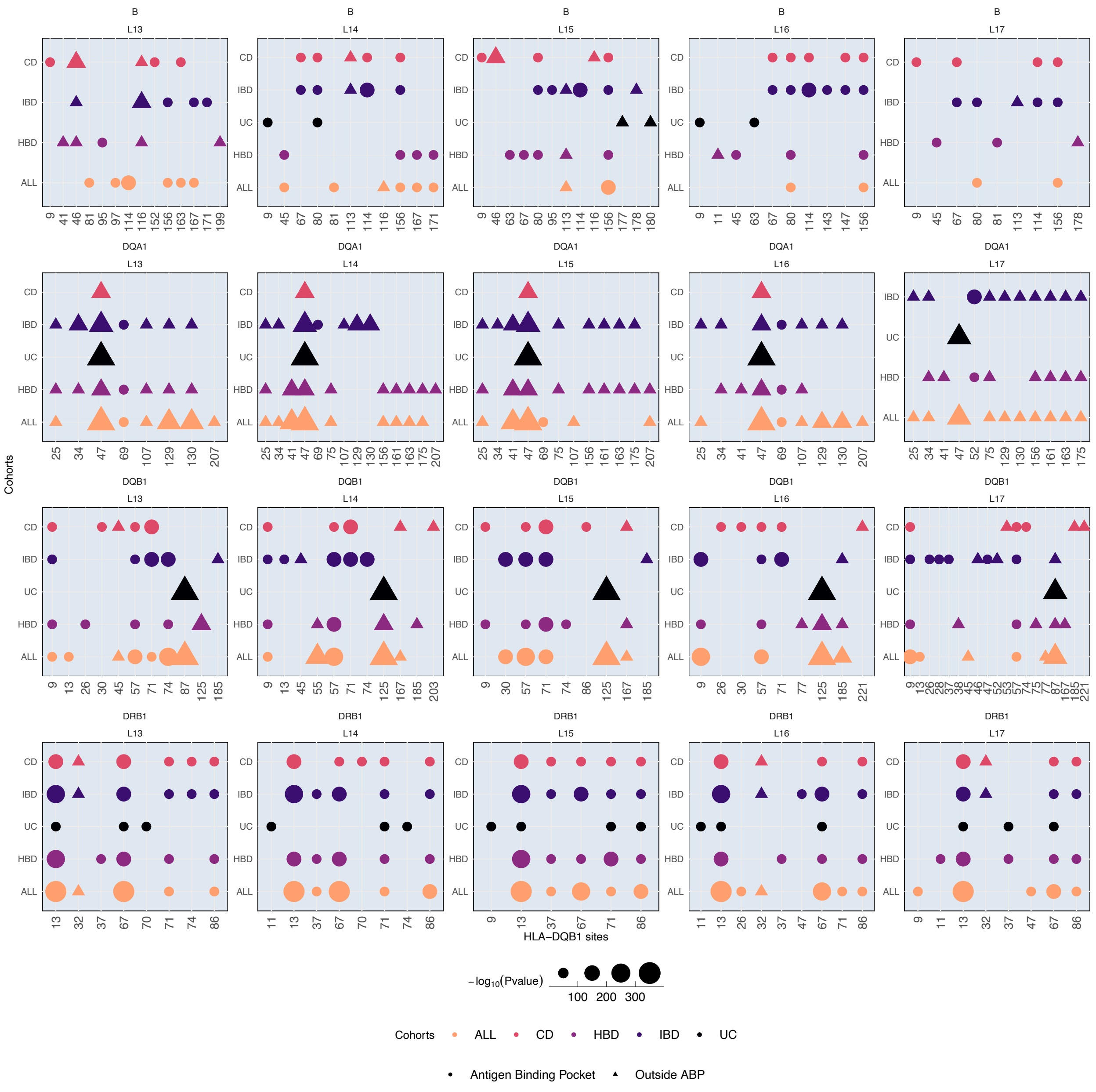


**Figure S15.** *The impact of HLA sites that are independent cdr3QTLs (X-axis) on the amino acid composition of the CDR3 loop, stratified by groups (CD, UC, IBD, HBD-healthy blood donors, ALL-combined datasets) and CDR3 length.* Each dot represents whether a specific HLA site was identified as *cdr3*QTL in a given dataset. The size of dots shows a statistical significance of *cdr3*QTL effects. The colour represents the dataset, and shape of a dot, either circle or triangle, indicate whether cdr3QTL is a peptide-binding pocket or outside.

#### S16: Structural analysis of *cdr3*QTLs in HLA-DRB1 and HLA-B

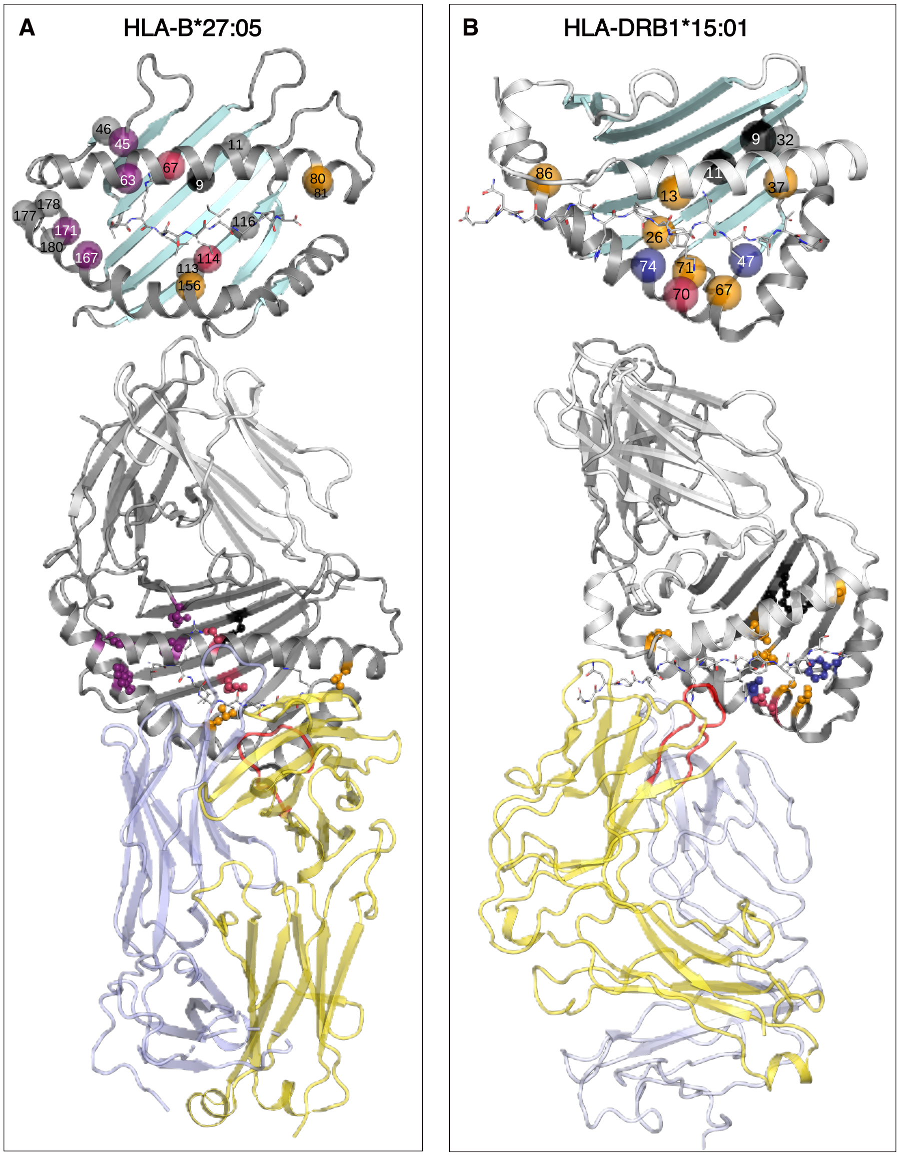


**Figure S16.** *Protein structural localisation of top cdr3QTLs in HLA-B and HLA-DR peptide-binding domains*. Top subplots with sites shown as C$\alpha$ spheres only, and HLA-peptide-TCR complexes at bottom subplots with sites “ball and stick” and colored as in **Figure 4A**. TCR alpha chains in light blue, TCR beta chains in yellow. The CDR3$\beta$ loop is highlighted in red. The peptide is depicted as sticks. **(A)** In HLA-B27 (PDB ID 7n2n), many *cdr3*QTL sites are buried in the peptide binding groove, either deeply on the central beta or on the flanking helices. In contact with the peptide N-terminus are 9, 45, 63, 67, 167, 171 pockets A and B. Sites located in the beta sheet located in central pockets B-E (114, 116, 156), and sites 80/81 in pocket F. Some *cdr3*QTL may interact with the TCR (63, 80, 167), or are neighbours of variant TCR contacting sites (67, 156, 171) *(5)*. **(B)** In HLA-DR15 (PDB ID 1ymm), most *cdr3*QTLs sites are buried in the peptide binding site, either deeply on the DRB1 central beta sheet or on the DRB1 helix. All sites except 9 and 32 contact the peptide directly (N-terminal P1: 86, central region P4-P7: 11, 13, 26, 67, 70, 71, C-terminal P9: 37). The flexible helical kink region 67-74, which was demonstrated to undergo structural movements during peptide processing and TCR binding, harbours several sites. Note that here an unconventional topology of an autoimmune TCR bound to DR15-MBP is shown, where the TCR is far from any cdr3QTL. In other DR structures, this region interacts with TCR. For example, in HLA-DR4 sites 67, 70 and 71 contact CDR3b 110/111 (PDB ID 8trl).

#### S17: Observed correlations between HLA alleles

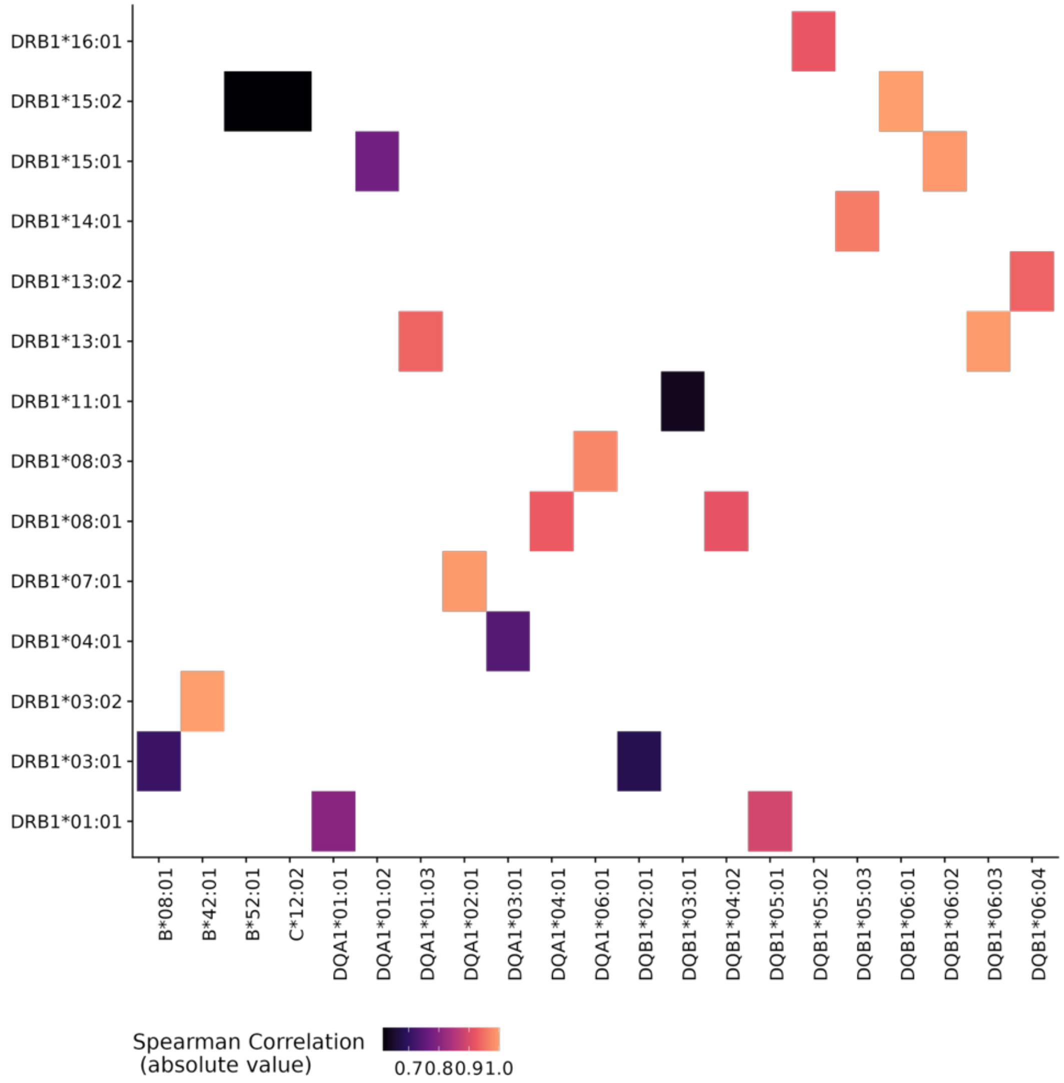


**Figure S17.** Spearman correlations between HLA class I and II alleles, observed in the combined dataset of HLA genotypes of healthy controls and individuals with IBD.

#### S18: Observed correlations between HLA sites

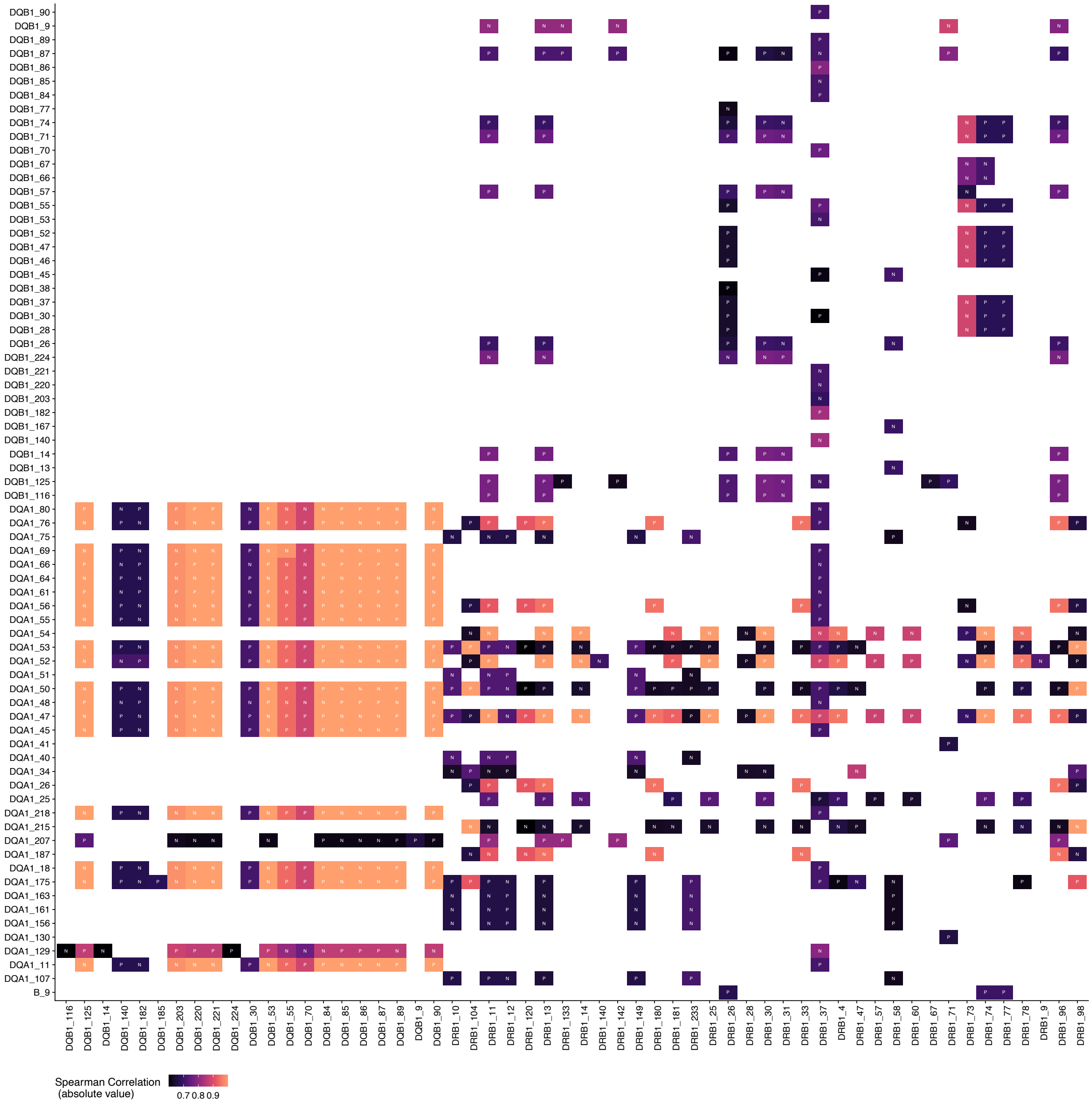


**Figure S18.** Spearman correlations between HLA class I and II residues, observed in the combined dataset of HLA genotypes of healthy controls and individuals with IBD. The values are in the absolute terms with the directionality coded by letters inside each cell, either P for positive and N for negative values.

#### S19: Conditional *cis*-haplotype analysis with PERMANOVA

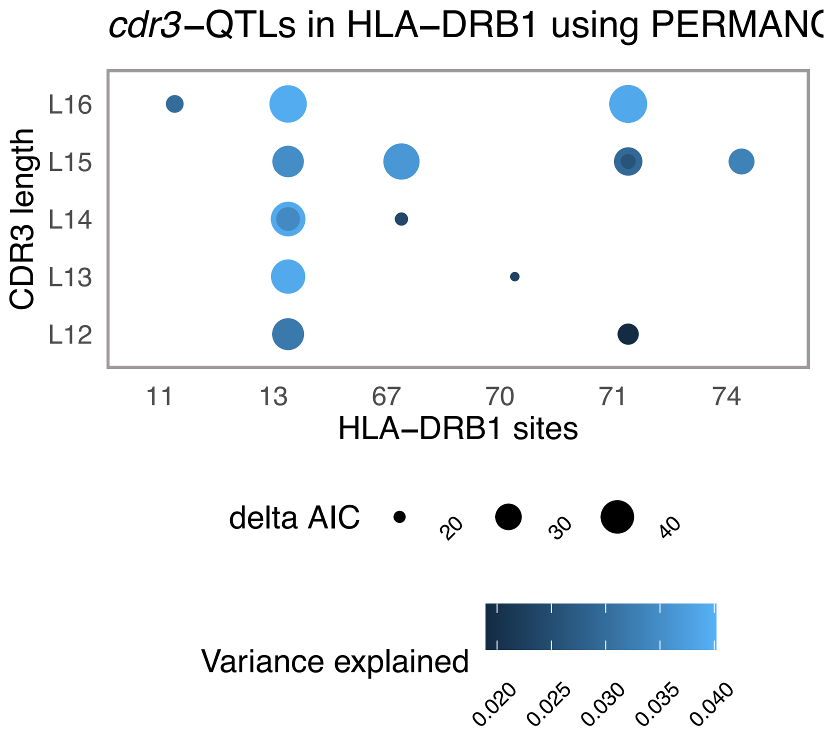


**Figure S19.** Validation of the key cdr3QTL signals in HLA-DRB1 with PERMANOVA. PERMANOVA accounts for the compositional nature of both HLA and TCR data, hence, provides much stronger certainty about HLA effects by supporting findings revealed by MANOVA.

#### S20: HLA-B alleles clustering based on *cdr3*QTLs

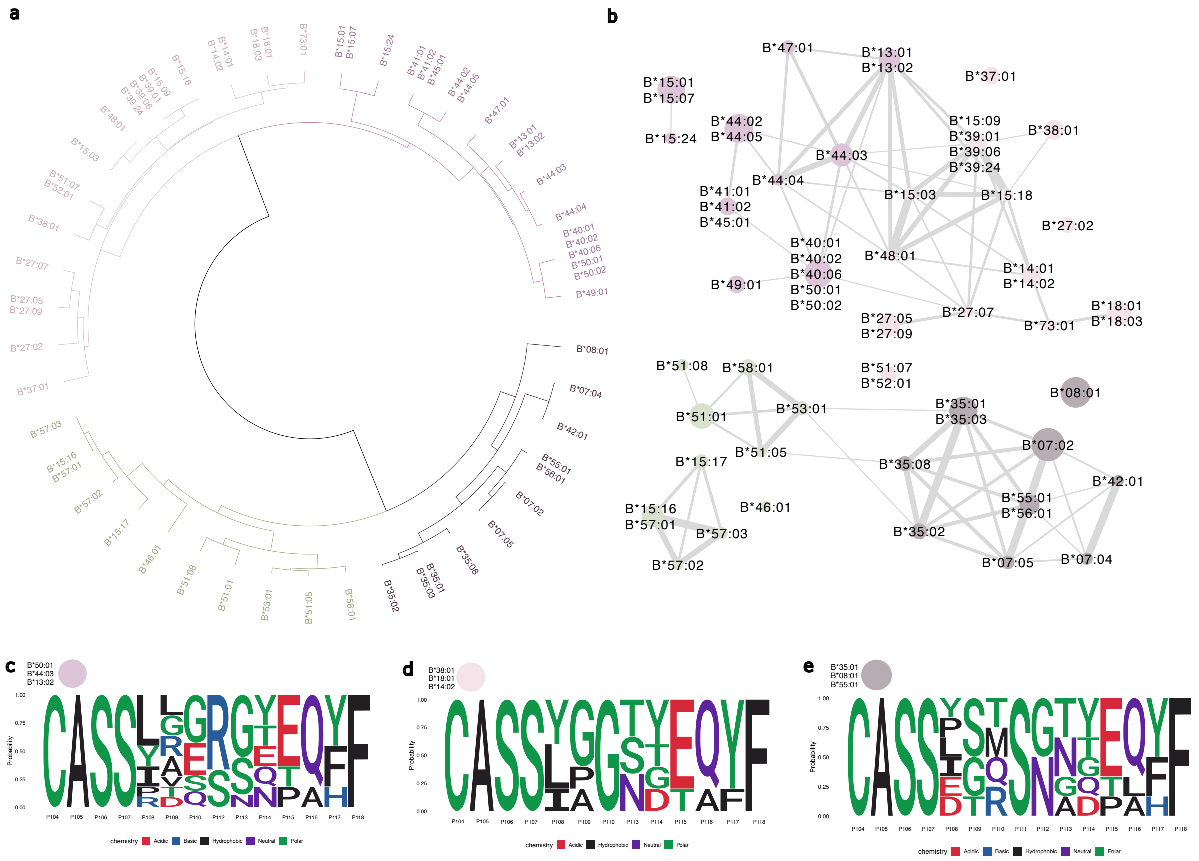


**Figure S20.** *Clustering of HLA-B alleles, based on physiochemical properties of amino acids, found at key cdr3QTLs with subsequent network analysis.* A: Common European population HLA alleles formed distinct clusters (color coded), coming from two main branches. B: The network analysis, based on amino acid physiochemical similarities at cdr3QTLs revealed that the key three clusters. C-E: Sequence logos illustrating the consensus amino acid motifs of public CDR3β clonotypes that demonstrated a significant statistical association with specific HLA alleles. Each logo represents the amino acid preferences (measured in probability, Y-axis) at each IMGT position within the CDR3 region (1-14 represent the IMGT positions, starting from P104 to P118). Clonotypes included in motif generation met an adjusted P-value after Bonferroni correction threshold of $p < 0.05$ from the association analysis. CDR3 amino acid motifs, defined but the public TRB clones, linked to any HLA-B alleles within each cluster. *The role of HLA-B alleles in chronic inflammatory diseases is less prominent than that of some HLA class II alleles. Nonetheless, we observed that signals within HLA-B were associated with the CDR3 composition of expanded singletons. HLA-B risk alleles for ankylosing spondylitis (HLA-B*27:05) (20,54), and type 1 diabetes (HLA-B*39:01/06 (20,55), HLA-B*18:01 and HLA-B*13:01/02 (20)) clustered tightly together, showing distinct effects on CDR3 linked to those alleles. We inferred CDR3 motifs from public TRB clones that were significantly linked to HLA-B alleles within each cluster. Compositions of CDR3 clones linked to one cluster were dissimilar from those, linked to another cluster. These motifs highlighted recurrent TCR patterns and HLA effects on the CDR3 loops, potentially mediated by antigens, as key cdr3QTLs localise to peptide-binding pockets.*

#### S21: Distribution of represented TCR binding epitope species in a public database VDJdb

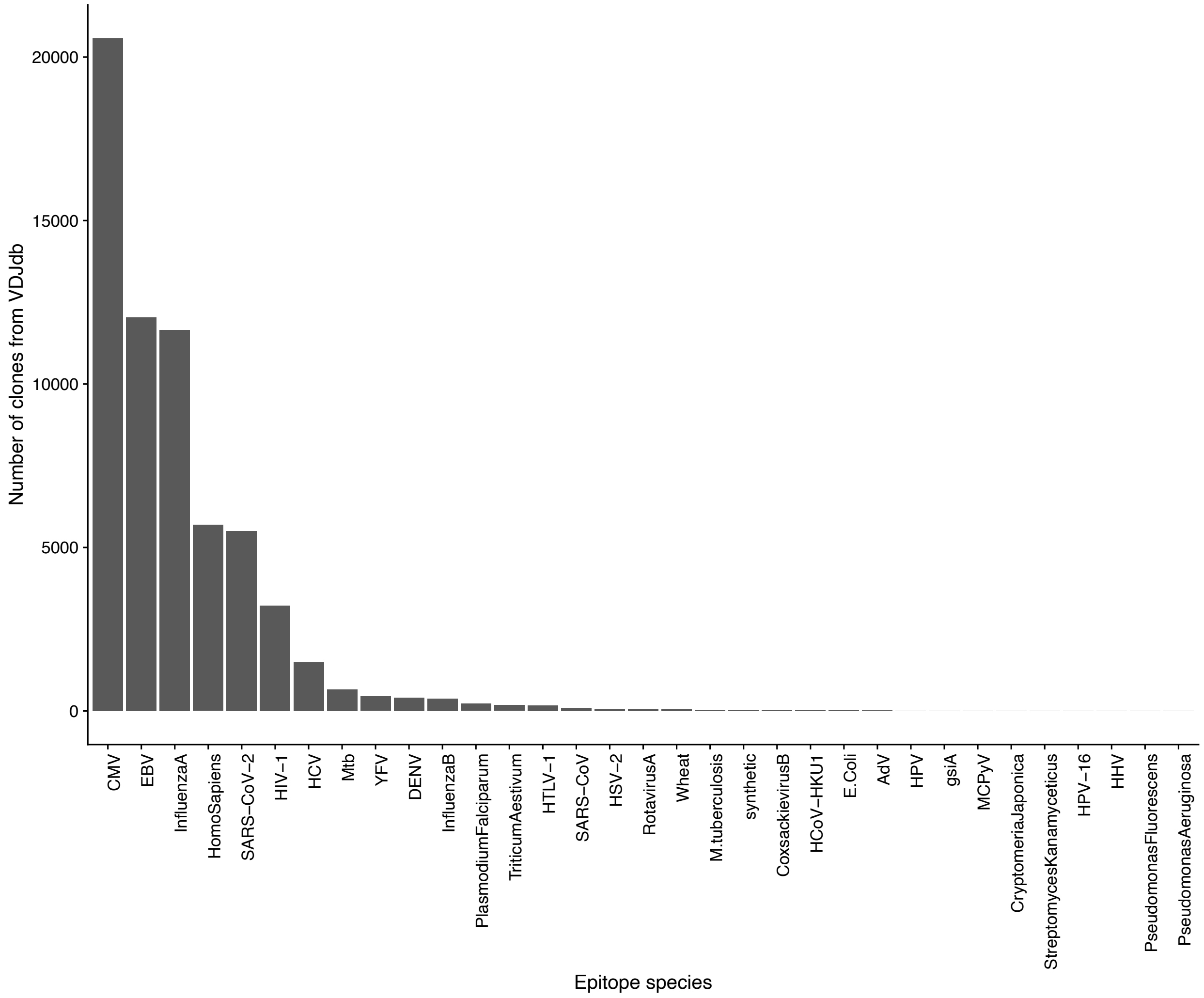


**Figure S21.** *The distribution of unique T-cell receptor (TCR) binding epitope species available in the VDJdb public database.* The X-axis represents the source species or pathogen of the epitopes, and the Y-axis shows the number of unique TCR-epitope pairs reported for each. The figure demonstrates a highly skewed distribution, with a significant majority of entries originating from a limited number of common pathogens, notably Cytomegalovirus (CMV), Epstein-Barr virus (EBV), and Influenza A. This highlights a substantial bias in the available public TCR-antigen data.

#### S22: Enrichment of influenced CDR3 amino acids by *cdr3*QTLs

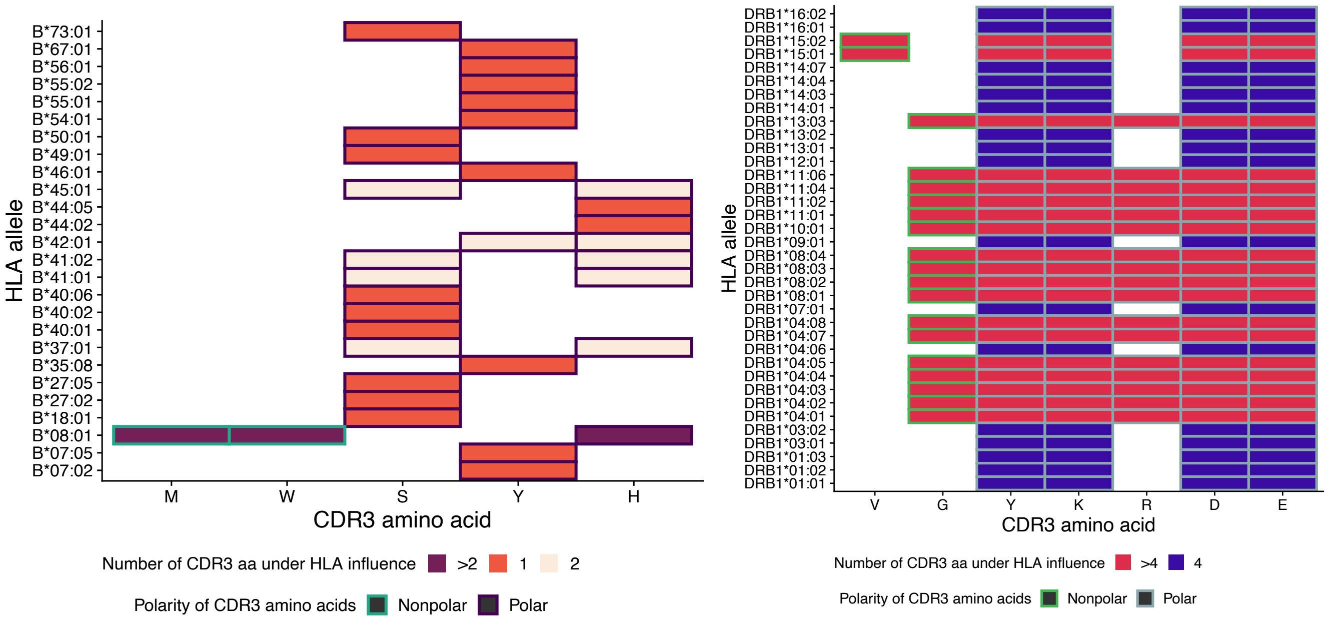


**Figure S22.** The enrichment of *cdr3*QTLs effects on specific amino acids across (X-axis) across different HLA-B and HLA-DRB1 alleles (Y-axis). The color intensity of each cell indicates binary state for each allele of how many CDR3 amino acid all site-specific *cdr3*QTL variants influence, red when more than 4, otherwise blue. The figure highlights that the risk alleles as DRB1*15:01, DRB1*04:01 for chronic inflammatory disease as MS, RA, T1D, SLE allele are enriched with the highest number of CDR3 amino acids under influence of their *cdr3*QTL variants, providing a mechanistic insight for the functional consequences for HLA risk alleles through their effects on CDR3.

#### S23: Structural analysis of IBD-specific cdr3QTLs in HLA-DQ

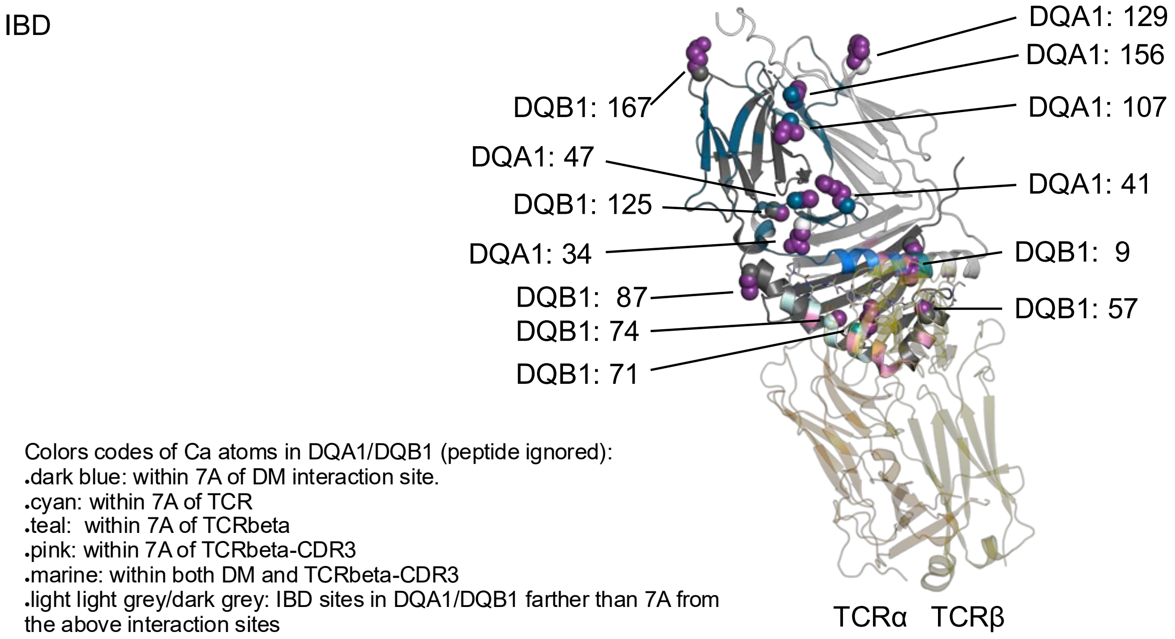


**Figure S22.** *Several DQA1 sites are positioned at or near regions known to interact with HLA-DM, as inferred from HLA-DM-DR protein structures.* These include DQA1:38 (41), 44 (47), 104 (107), 126 (129), and 153 (156). Among these, DQA1:44 (47) and 104 (107), along with DQA1:31 (34) and 153 (156), also reside at HLA chain interfaces. Similarly, DQB1:9, 57, and 125 engage in interactions with the DQA chain. These polymorphic sites are often structural neighbors of conserved residues crucial for HLA-DM-HLA-II and HLA interchain interactions. Localization of DQA1-DQB1 cdr3QTLs in the protein structure of DQ2.5/gliadin peptide/TCR (PDB ID 4ozf). At or near the PBS, buried are DQA1:31(34), DQB1:9 and DQB1:57, while DQB1:87 points to the outside of the groove and DQB1:71 and DQB1:74 on the PBS DQB1 helix, In direct peptide contact are in P9 DQB1:9 and DQB1:57, while in P4-7 DQB1:71 and P4 DQB1:74. The other cdr3QTLs locate at or near regions that are predicted to interact with HLA-DM (38(41), 44(47), 104(107), 126(129), 153(156)) or are at or near interacting sites for the respective other HLA chain (DQA1:44 (47), 104 (107), 31 (34), 153 (156). DQB1:9, 57, 125). Further, DQB1:167 locates on a membrane-proximal loop and DQA1:207 within the transmembrane helix (not shown). TCRa in orange, TCRb in yellow. A cutoff of 7A was chosen to define direct or indirect interacting residues. Following, DQA1/DQB1 sites near HLA-DM are colored dark blue. Sites near the TCR/TCRb are colored in cyan/teal and those near the CDR3β only in pink. In marine, the region in vicinity of both, CDR3β and HLA-DM.

#### Supplemental Tables

**Table S1: Public TRB clones with a strong association with common HLA alleles**

| HLA allele | TRB clone | P_adj(Bonf) |
|---|---|---|
| B*08:01 | CASSDSTSGTDTQYF_TCRBV06-04_TCRBJ02-03 | 0.00441830169314368 |
| B*13:02 | CASSLAGRGETQYF_TCRBV07-02_TCRBJ02-05 | 3.69648183337154e-10 |
| B*13:02 | CASSRDSSYEQYF_TCRBV27-01_TCRBJ02-07 | 8.44776129034899e-09 |
| B*13:02 | CASSITGSSYEQYF_TCRBV19-01_TCRBJ02-07 | 5.87674147688693e-06 |
| B*13:02 | CASSYREGQPQHF_TCRBV06-05_TCRBJ01-05 | 9.47631239932666e-05 |
| B*13:02 | CASSLLNEQFF_TCRBV07-02_TCRBJ02-01 | 8.91387487356174e-05 |
| B*14:02 | CASSYAYEQYF_TCRBV06-02_TCRBJ02-07 | 0.00382726362275217 |
| B*14:02 | CASSYGGEQYF_TCRBV06-05_TCRBJ02-07 | 0.0119100109524226 |
| B*14:02 | CASSLGNTEAFF_TCRBV05-05_TCRBJ01-01 | 0.00483033965918496 |
| B*14:02 | CASSPGYEQYF_TCRBV05-01_TCRBJ02-07 | 0.000121509668448661 |
| B*15:01 | CASSLEGQGTDTQYF_TCRBV05-01_TCRBJ02-03 | 6.728260523616e-08 |
| B*18:01 | CASSLPGSYEQYF_TCRBV07-09_TCRBJ02-07 | 2.39728540255174e-05 |
| B*18:01 | CASSPTGSTDTQYF_TCRBV18-01_TCRBJ02-03 | 0.00610357656223137 |
| B*27:05 | CASSPGTGGNQPQHF_TCRBV27-01_TCRBJ01-05 | 0.00706927848935245 |
| B*35:01 | CASSISRNTGELFF_TCRBV19-01_TCRBJ02-02 | 2.11129703714985e-05 |
| B*35:01 | CASSEGMNTEAFF_TCRBV02-01_TCRBJ01-01 | 8.61847612014642e-17 |
| B*35:01 | CASSLGQAYEQYF_TCRBV27-01_TCRBJ02-07 | 0.0180546900610274 |
| B*38:01 | CASSIGTDTQYF_TCRBV19-01_TCRBJ02-03 | 0.000191922675884712 |
| B*44:03 | CASSIGSTEAFF_TCRBV19-01_TCRBJ01-01 | 0.000806193992600912 |
| B*44:03 | CASSPVNTEAFF_TCRBV18-01_TCRBJ01-01 | 0.00121560979087621 |
| B*44:03 | CASSLAGRGETQYF_TCRBV07-02_TCRBJ02-05 | 2.09850509538397e-05 |
| B*44:03 | CASSYREGQPQHF_TCRBV06-05_TCRBJ01-05 | 1.5289055883305e-05 |
| B*44:03 | CASSLLNEQFF_TCRBV07-02_TCRBJ02-01 | 0.00123416050628501 |
| B*50:01 | CASSLRGTEAFF_TCRBV05-06_TCRBJ01-01 | 0.00172305325808653 |
| B*55:01 | CASSYSNQPQHF_TCRBV06-02_TCRBJ01-05 | 9.86574659080646e-11 |
| DRB1*01:01 | CASSPGQGPGNTIYF_TCRBV11-03_TCRBJ01-03 | 9.27774297733488e-16 |
| DRB1*01:01 | CSARVAGGTDTQYF_TCRBV20_TCRBJ02-03 | 0.0158219241855209 |
| DRB1*01:02 | CASSIGGGQETQYF_TCRBV19-01_TCRBJ02-05 | 0.0199883786354578 |
| DRB1*03:01 | CASSDSTSGTDTQYF_TCRBV06-04_TCRBJ02-03 | 1.22079787501096e-12 |
| DRB1*03:01 | CASSDSTGGTDTQYF_TCRBV06-04_TCRBJ02-03 | 0.00732292779596251 |
| DRB1*04:01 | CASSDSTSGTDTQYF_TCRBV06-04_TCRBJ02-03 | 2.45928309986289e-05 |
| DRB1*04:01 | CASSGTSTDTQYF_TCRBV06-04_TCRBJ02-03 | 0.0340788151657615 |
| DRB1*04:01 | CASSDRNTGELFF_TCRBV06-04_TCRBJ02-02 | 1.06344360646189e-05 |
| DRB1*04:01 | CASSLRGETQYF_TCRBV05-01_TCRBJ02-05 | 0.0086992197107127 |
| DRB1*04:03 | CASSFTGGNQPQHF_TCRBV12_TCRBJ01-05 | 0.000115374730073316 |
| DRB1*04:03 | CASSDSNTGELFF_TCRBV06-04_TCRBJ02-02 | 0.000798161550535289 |
| DRB1*04:03 | CASSPGQGNQPQHF_TCRBV28-01_TCRBJ01-05 | 0.0496384974175726 |
| DRB1*04:03 | CASSLVVNTEAFF_TCRBV07-09_TCRBJ01-01 | 0.00465350457242441 |
| DRB1*04:03 | CASSLQGNTEAFF_TCRBV19-01_TCRBJ01-01 | 0.00101267689544924 |
| DRB1*04:03 | CASSYSGANVLTF_TCRBV06-02_TCRBJ02-06 | 0.00458108045035271 |
| DRB1*04:03 | CASSGTGGNQPQHF_TCRBV19-01_TCRBJ01-05 | 0.000120160801694751 |
| DRB1*04:03 | CASSLSDYGYTF_TCRBV28-01_TCRBJ01-02 | 0.00032071607341711 |
| DRB1*04:03 | CAWSVQGDQPQHF_TCRBV30-01_TCRBJ01-05 | 0.025977255108908 |
| DRB1*07:01 | CAWSRDSGSGNTIYF_TCRBV30-01_TCRBJ01-03 | 0.0146403024924402 |
| DRB1*08:01 | CASSPRGDTEAFF_TCRBV18-01_TCRBJ01-01 | 0.00172738048090221 |
| DRB1*08:01 | CSARRTNTEAFF_TCRBV20_TCRBJ01-01 | 0.022630769512855 |
| DRB1*09:01 | CASSVGDYNEQFF_TCRBV09-01_TCRBJ02-01 | 0.00191811816788478 |
| DRB1*09:01 | CASSPGTGDEKLFF_TCRBV18-01_TCRBJ01-04 | 4.88110396117025e-11 |

| | | |
|---|---|---|
| DRB1*11:01 | CASSVGQGSYNEQFF_TCRBV09-01_TCRBJ02-01 | 4.18786842584594e-10 |
| DRB1*11:01 | CASSQAGMNTEAFF_TCRBV14-01_TCRBJ01-01 | 0.0474040008986939 |
| DRB1*11:04 | CASSPSTDTQYF_TCRBV11-02_TCRBJ02-03 | 0.0032194220536275 |
| DRB1*11:04 | CASSFGQGGEKLFF_TCRBV05-01_TCRBJ01-04 | 3.21185771647602e-05 |
| DRB1*11:04 | CASSPSYEQYF_TCRBV11-02_TCRBJ02-07 | 0.00164245444058575 |
| DRB1*11:04 | CASSYLQGNTEAFF_TCRBV06-05_TCRBJ01-01 | 1.63829524412395e-09 |
| DRB1*12:01 | CASSLSGGNYGYTF_TCRBV27-01_TCRBJ01-02 | 0.0206202834858767 |
| DRB1*13:01 | CASSPGQGPGNTIYF_TCRBV11-03_TCRBJ01-03 | 0.0146673884607074 |
| DRB1*13:01 | CASSLGGYEQYF_TCRBV11-02_TCRBJ02-07 | 0.0055613298682754 |
| DRB1*13:01 | CASSLRETQYF_TCRBV07-09_TCRBJ02-05 | 0.00010979676027439 |
| DRB1*13:01 | CAWSVRGNQPQHF_TCRBV30-01_TCRBJ01-05 | 1.59276986921251e-16 |
| DRB1*13:01 | CASRRNTEAFF_TCRBV06-05_TCRBJ01-01 | 0.000723754880570611 |
| DRB1*13:01 | CASSLAGTGGGYTF_TCRBV05-01_TCRBJ01-02 | 1.09420596619986e-08 |
| DRB1*13:01 | CASSPPGGSYEQYF_TCRBV18-01_TCRBJ02-07 | 0.000112421866420299 |
| DRB1*13:01 | CSARGAGNTIYF_TCRBV20_TCRBJ01-03 | 0.0418527925400065 |
| DRB1*13:01 | CASSGQPNTEAFF_TCRBV02-01_TCRBJ01-01 | 0.00719966261469351 |
| DRB1*13:02 | CAWSRDSGSGNTIYF_TCRBV30-01_TCRBJ01-03 | 6.86898054242068e-08 |
| DRB1*13:02 | CASSRTYEQYF_TCRBV06-02_TCRBJ02-07 | 0.00440817252252118 |
| DRB1*13:03 | CASSPGQGNYGYTF_TCRBV03-01_TCRBJ01-02 | 2.52702038713625e-06 |
| DRB1*13:03 | CASSAGYEQYF_TCRBV02-01_TCRBJ02-07 | 3.59156891772052e-05 |
| DRB1*13:03 | CSAGQGSYEQYF_TCRBV20_TCRBJ02-07 | 0.0057486021491361 |
| DRB1*13:03 | CSARVGGNQPQHF_TCRBV20_TCRBJ01-05 | 0.00602925724141603 |
| DRB1*13:03 | CSARGQLNTEAFF_TCRBV20_TCRBJ01-01 | 0.0481591141552195 |
| DRB1*13:03 | CASSLGQTYNEQFF_TCRBV05-01_TCRBJ02-01 | 3.78061038965515e-05 |
| DRB1*13:03 | CASSPDRVYGYTF_TCRBV18-01_TCRBJ01-02 | 0.00819870703391377 |
| DRB1*13:03 | CASSLRHLNTEAFF_TCRBV12_TCRBJ01-01 | 0.00130455957276149 |
| DRB1*13:03 | CATSRDPGGNQPQHF_TCRBV15-01_TCRBJ01-05 | 0.00151072580577681 |
| DRB1*14:01 | CASSLQGSNQPQHF_TCRBV12_TCRBJ01-05 | 0.0242182006644294 |
| DRB1*14:01 | CASSYTGELFF_TCRBV12-03_TCRBJ02-02 | 0.00432823579807005 |
| DRB1*14:01 | CSASNTEAFF_TCRBV20-01_TCRBJ01-01 | 0.0194348619321201 |
| DRB1*14:01 | CSARTTYEQYF_TCRBV20-01_TCRBJ02-07 | 1.0391917585925e-05 |
| DRB1*14:01 | CASSLSGGYEQYF_TCRBV07-09_TCRBJ02-07 | 0.0410837382371977 |
| DRB1*15:01 | CASSQAGMNTEAFF_TCRBV14-01_TCRBJ01-01 | 0.0124935191411906 |
| DRB1*15:02 | CASSLDGYSNQPQHF_TCRBV05-01_TCRBJ01-05 | 0.00546148412807793 |
| DRB1*16:01 | CASSDSSTDTQYF_TCRBV06-04_TCRBJ02-03 | 0.000264670848810306 |
| DRB1*16:01 | CASSPGQGNQPQHF_TCRBV12_TCRBJ01-05 | 1.40962177817232e-07 |
| DRB1*16:01 | CASSDSSGGADTQYF_TCRBV06-04_TCRBJ02-03 | 0.0054435248648516 |
| DRB1*16:01 | CASSLKGNTEAFF_TCRBV12-03_TCRBJ01-01 | 6.9356416407486e-05 |
| DRB1*16:01 | CASSDRDTDTQYF_TCRBV06-04_TCRBJ02-03 | 0.0149448302404517 |
| DRB1*16:01 | CASSDSGTDTQYF_TCRBV06-04_TCRBJ02-03 | 0.00618262629054094 |
| DRB1*16:01 | CASSDSTGGADTQYF_TCRBV06-04_TCRBJ02-03 | 0.0458944728128265 |

**Table S2: Annotated with peptide-binding pockets resulting HLA-CDR3 associations from CDR3-QTL discovery analysis.**

CDR3-QTL-in-IBD-Lokes-et-al-2025.csv
